# Multiscale integration organizes hierarchical computation in human auditory cortex

**DOI:** 10.1101/2020.09.30.321687

**Authors:** Sam V Norman-Haignere, Laura K. Long, Orrin Devinsky, Werner Doyle, Ifeoma Irobunda, Edward M. Merricks, Neil A. Feldstein, Guy M. McKhann, Catherine A. Schevon, Adeen Flinker, Nima Mesgarani

## Abstract

To derive meaning from sound, the brain must integrate information across tens (e.g. phonemes) to hundreds (e.g. words) of milliseconds, but the neural computations that enable multiscale integration remain unclear. Prior evidence suggests that human auditory cortex analyzes sound using both generic acoustic features (e.g. spectrotemporal modulation) and category-specific computations, but how these putatively distinct computations integrate temporal information is unknown. To answer this question, we developed a novel method to estimate neural integration periods and applied the method to intracranial recordings from human epilepsy patients. We show that integration periods increase three-fold as one ascends the auditory cortical hierarchy. Moreover, we find that electrodes with short integration periods (~50-150 ms) respond selectively to spectrotemporal modulations, while electrodes with long integration periods (~200-300 ms) show prominent selectivity for sound categories such as speech and music. These findings reveal how multiscale temporal analysis organizes hierarchical computation in human auditory cortex.

---

Time is the fundamental dimension of sound, and temporal integration is thus fundamental to audition. To recognize a complex structure like a word, the brain must integrate information across a wide range of timescales from tens (e.g. phonemes) to hundreds (e.g. syllables) of milliseconds (**Fig S1**)^1^. But how human auditory cortex accomplishes this feat is unclear.

One prominent hypothesis posits that short and long-term temporal structure are analyzed asymmetrically across the two hemispheres, with the left hemisphere integrating over short timescales, and the right hemisphere integrating over long timescales^2–4^. Another influential hypothesis is that the auditory cortex integrates across time hierarchically, with short-term structure analyzed bilaterally in primary auditory cortex and longer-term structure analyzed in non-primary regions^5–7^. This question remains unresolved, despite intensive debate over two decades, because the integration period of human cortical regions is unknown.

Understanding temporal integration is critical for understanding how important sound categories like speech and music are processed^2,6,8^. While prior studies have revealed non-primary neural populations selective for speech and music^9–13^, little is known about how these neural populations integrate information in speech and music. One possibility is that category-selective neural populations integrate over many timescales in order to code category-specific structure at short^14,15^ (e.g. phonemes) and long^8^ timescales (e.g. syllables and words; **Fig S1**). Alternatively, short-term structure might be analyzed by general-purpose acoustic representations in primary auditory cortex^16^ and then integrated over long timescales to form category-specific neural representations in non-primary regions.

Here, we test these hypotheses by developing a novel method for measuring neural integration periods. Integration periods are often defined as the time window when stimuli alter the neural response^17,18^. Although this definition is simple and general, there is no simple and general method to estimate integration periods. Many methods exist for inferring linear integration periods with respect to a spectrogram^15,19–21^, but human cortical responses exhibit prominent nonlinearities particularly in non-primary regions. Flexible, nonlinear models are challenging to fit given limited neural data^20,22^, and even if one succeeds, it is not obvious how to measure the model’s integration period. Methods for assessing temporal modulation selectivity^6,23,24^ are insufficient, since a neuron could respond to fast modulations over a long window or to a complex structure like a word that is poorly described by its modulation content. Finally, temporal scrambling can reveal selectivity for naturalistic temporal structure^12,18,25^, but many regions in auditory cortex show no difference between intact and scrambled sounds.

To overcome these limitations, we developed a method that directly estimates the time window when stimuli alter a neural response (the temporal context invariance or TCI paradigm; **Fig 1**). We present sequences of natural stimuli in a random order such that the same segment occurs in different contexts. While context has many meanings^26^, here we simply define context as the stimuli which surround a segment. If the integration period is shorter than the segment duration, there will be a moment when it is fully contained within each segment. As a consequence, the response to each segment will be unaffected by the surrounding segments. We can therefore estimate the integration period by determining the minimum segment duration needed to achieve a context invariant response.

**Fig 1.**
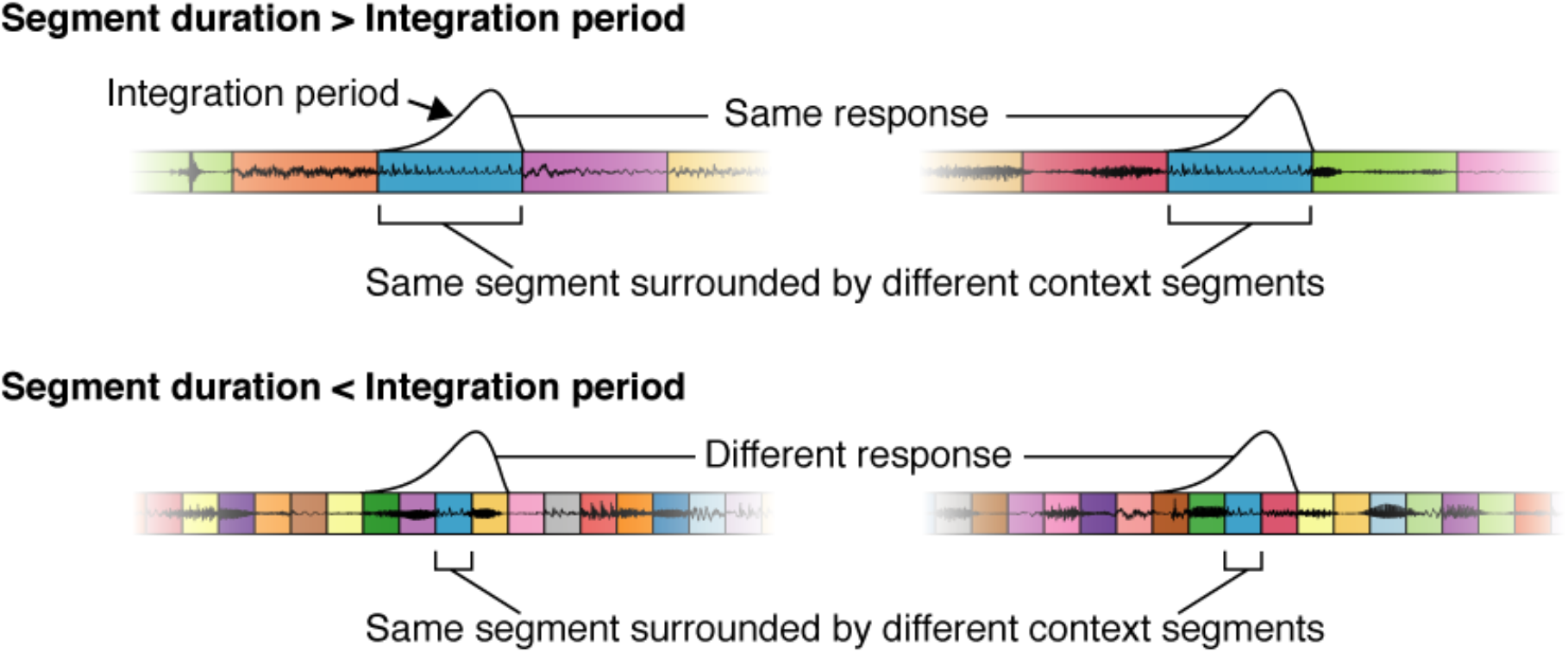
Temporal context invariance (TCI) paradigm. Schematic of the paradigm used to measure integration periods. Segments of natural stimuli are presented using two different random orderings. As a consequence, the same segment occurs in two different contexts (different surrounding segments). If the segment duration is longer than the integration period (top panel), there will be a moment when the integration period is fully contained within each segment. As a consequence, the response at that moment will be unaffected by the surrounding context segments. If the segment duration is shorter than the integration period (bottom panel), the integration period will always overlap the surrounding context segments, and they can therefore alter the response. The goal of the TCI paradigm is to estimate the minimum segment duration needed to achieve a context invariant response. This figure plots waveforms for an example sequence of segments that share the same central segment. Segment boundaries are demarcated by colored boxes. The hypothesized integration period is plotted above each sequence at the moment when it best overlaps the shared segment.

TCI does not make any assumptions about the type of response being measured. As a consequence, the method is applicable to sensory responses from any modality, stimulus set, or recording method. We applied TCI to intracranial EEG (iEEG) recordings collected from patients undergoing surgery for intractable epilepsy. Such recordings provide a rare opportunity to measure human brain responses with spatiotemporal precision, which is essential to studying temporal integration. We used a combination of depth and surface electrodes to record from both primary regions in the lateral sulcus as well as non-primary regions in the superior temporal gyrus (STG), unlike many iEEG studies that have focused on just the lateral sulcus^27^ or STG^15,19^. The precision and coverage of our recordings was essential to revealing how the human auditory cortex integrates across multiple timescales.

## Results

We recorded iEEG responses to sequences of natural sound segments that varied in duration (from 31 ms to 2 sec in octave steps). For each segment duration, we created two 20-second sequences, each with a different random ordering of the same segments (concatenated using cross-fading to avoid boundary artifacts). Segments were excerpted from 10 natural sounds (**Table S1**), selected to be diverse so they differentially drive responses throughout auditory cortex. The same natural sounds were used for all segment durations, which limited the number of sounds we could test given the limited time with each patient; but our key results were robust across the sounds tested (see *Anatomical organization* for the results of all robustness analyses). Because our goal was to characterize integration periods during natural listening, we did not give subjects a formal task. To encourage subjects to listen to the sounds, we asked them to occasionally rate how scrambled the last stimulus sequence was (shorter segment durations sound more scrambled; if patients were in pain or confused we simply asked them to listen).

### Assessing context invariance via the cross-context correlation

We measured the broadband gamma power of each electrode to each sound sequence, which is thought to approximately reflect aggregate neural activity in a local region^28,29^ (70-140 Hz; results were robust to the frequency range used). For each electrode, we aligned its response to all segments of a given duration in a matrix, which we refer to as the segment-aligned response (SIR) matrix (**Fig 2a**). Each row of the SIR matrix contained the response timecourse surrounding a single segment, aligned to segment onset. Different rows thus correspond to different segments and different columns correspond to different lags relative to segment onset. We computed two versions of the SIR matrix using the two different contexts for each segment, extracted from the two different sequences. The central segment was the same between the contexts, but the surrounding segments were different.

**Fig 2.**
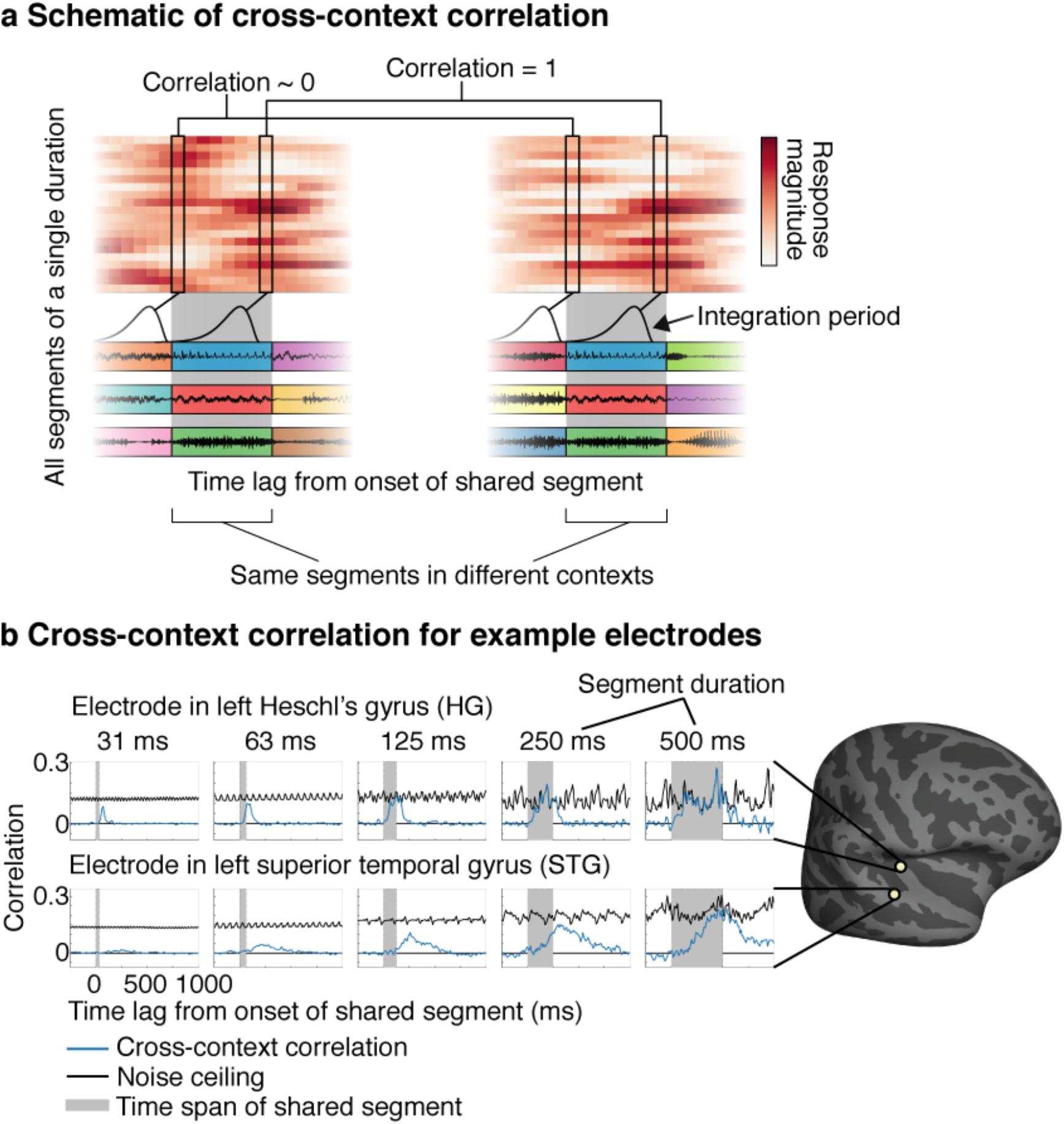
Cross-context correlation. **a,** Schematic of the analysis used to assess context invariance for a single electrode and segment duration. See text for description. **b,** The cross-context correlation (blue line) and noise ceiling (black line) are shown for two example electrodes from the left hemisphere of the same patient, one in Heschl’s gyrus (HG, top panel) and one in the superior temporal gyrus (STG, bottom panel). Each plot shows a different segment duration. The gray region shows the time interval when the shared segment was present (i.e. the gray region in panel a). The STG electrode required longer segment durations for the cross-context correlation to reach the noise ceiling, and the build-up of the cross-context correlation with lag was slower for the STG electrode.

Our goal was to assess if there was a lag when the response was the same across contexts. We instantiated this idea by correlating corresponding columns across SIR matrices from different contexts (the “cross-context correlation”, schematized in **Fig 2a**). At segment onset (lag=0), the cross-context correlation should be near zero, since the integration period must overlap the preceding segments, which were random across contexts. As time progresses, the integration period should start to overlap the shared segment, and the cross-context correlation should increase. Critically, if the integration period is less than the segment duration, there should be a lag where the integration period is fully contained within the shared segment, and the response should thus be the same, yielding a correlation of 1 modulo noise. To correct for noise, we measured the test-retest correlation when the context was the same, which provides a noise ceiling for the cross-context correlation.

The shorter segments tested in our study were created by subdividing the longer segments. As a consequence, we could also consider cases where a segment was a subset of a longer segment and thus surrounded by its natural context, in addition to the case described so far when a segment is surrounded by random other segments. Since our analysis requires that the two contexts being compared are different, one of the two contexts must always be random, but the other context can be random or natural. In practice, we found similar results using random and natural contexts, and thus pooled across both types of context for maximal statistical power (see *Anatomical organization* for results comparing random and natural contexts).

We plot the cross-context and noise ceiling for segments of increasing duration for two example electrodes from the same subject: an electrode in left posteromedial Heschl’s gyrus (HG) and one in the left superior temporal gyrus (STG) (**Fig 2b**). The periodic variation in the noise ceiling is an inevitable consequence of correlating across a fixed set of segments (see *Cross-context correlation* in the Methods for an explanation). For the HG electrode, the cross-context correlation started at zero and quickly rose. Critically, for segment durations greater than or equal to 125 milliseconds, there was a lag where the cross-context correlation equaled the noise ceiling, indicating a context invariant response. For longer segments (250 or 500 ms), the cross-context correlation remained yoked to the noise ceiling for an extended duration indicating that the integration period remained within the shared segment for an extended time period. This pattern is what one would expect for an integration period that is ~125 milliseconds, since stimuli falling outside of this window have little effect on the response.

By comparison, the results for the STG electrode suggest a much longer integration period. Only for segment durations of approximately 500 milliseconds did the cross-context correlation approach the noise ceiling, and its build-up and fall-off with lag was considerably slower. This pattern is what one would expect for a longer integration period, since it takes more time for the integration period to fully enter and exit the shared segment. Virtually all electrodes with a reliable response to sound exhibited a similar pattern, but the segment duration and lag needed to achieve an invariant response varied substantially (**Fig S2** shows 20 representative electrodes). This observation indicates that auditory cortical responses have a meaningful integration period, outside of which responses are largely invariant, but the extent of this integration period varies across the cortex.

### Model-estimated integration periods

In theory, one could estimate the integration period extent as the shortest segment duration for which the peak of the cross-context correlation exceeds some fraction of the noise ceiling. This approach, however, would be noise-prone since a single noisy data point at one lag and segment duration could alter the estimated integration period. To overcome this issue, we used a model to infer the integration period that best-predicted the cross-context correlation across all lags and segment durations.

We modeled temporal integration periods using a Gamma-distributed window, which is a standard, unimodal distribution commonly used to model integration periods (**Fig 3a**)^30^. We varied the width and center of the model integration period, excluding combinations of widths and centers that resulted in a non-causal window since this would imply the response depends upon future stimuli. The width of the integration period is the key parameter we would like to estimate, and was defined as the smallest interval that contained 75% of the window’s mass. The center of the integration period was defined as the window’s median and reflects the overall delay between the integration period and the response. We also varied the window shape from more exponential to more bell-shaped, but found the shape had little influence on the results (see *Anatomical organization*).

**Fig 3.**
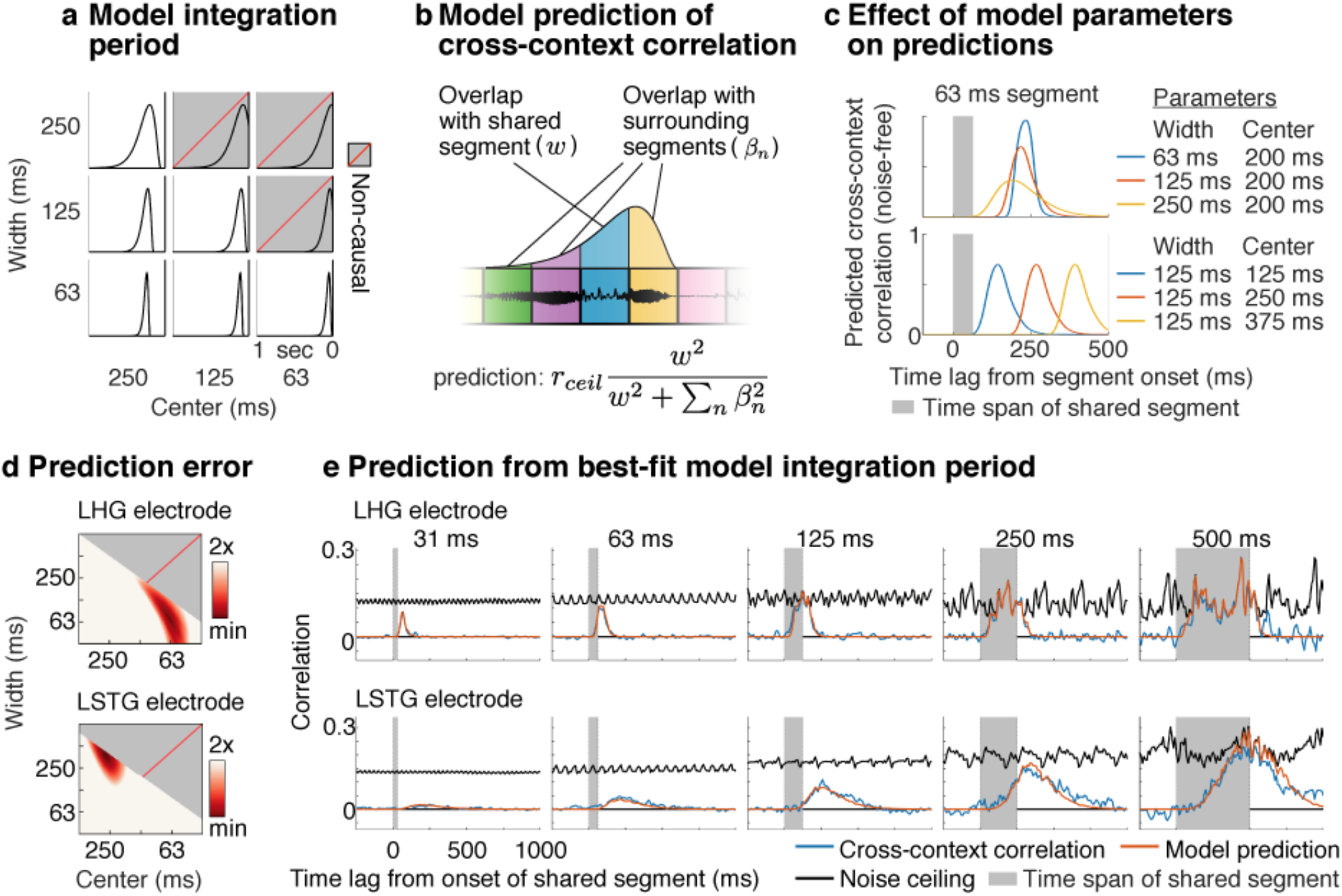
Model-estimated integration periods. **a**, Temporal integration periods were modeled using a Gamma-distributed window. The width and center of the model integration period were varied, excluding combinations of widths and centers that resulted in a non-causal window (gray boxes with dashed red line). **b,** Schematic showing how the cross-context correlation was predicted from the model integration period. For each segment duration and lag, we measured how much the integration period overlapped the shared central segment (*w*, blue segment) vs. all surrounding context segments (*β*_*n*_, yellow, purple, and green segments). The cross-context correlation should reflect the fraction of the response variance due to the shared segment, multiplied by the noise ceiling (*r*_*ceil*_). The variance due to each segment is given by the squared overlap with the model integration period. **c,** Illustration of how the integration period width (top panel) and center (bottom panel) alter the model’s prediction for a single segment duration (63 milliseconds). Increasing the width lowers and stretches-out the predicted cross-context correlation, while increasing the center shifts the cross-context correlation to later lags. **d**, The prediction error for model windows of varying widths and centers for the example electrodes from **Figure 2b**. Redder colors indicate lower error. **e**, The measured and predicted cross-context correlation for the best-fit integration period with lowest error (same format as **Fig 2b**).

The cross-context correlation depends on the degree to which the integration period overlaps the shared segment vs. the surrounding context segments (**Fig 2a**). We therefore predicted the cross-context correlation by measuring the overlap between the model integration period and each segment, separately for all lags and segment durations (**Fig 3b**). The equation used to predict the cross-context correlation from these overlap measures is shown in **Figure 3b** and described in the legend. A formal derivation is given in the Methods.

**Figure 3c** illustrates how changing the width and center of the model integration period alters the predicted correlation. Increasing the width lowers the peak of the cross-context correlation, since a smaller fraction of the integration period overlaps the shared segment at the moment of maximum overlap. The build-up and fall-off with lag is also more gradual for wider integration periods since it takes longer for the integration period to enter and exit the shared segment. Increasing the center simply shifts the cross-context correlation to later lags, since the delay is longer, but the width is unchanged.

We varied the model parameters and calculated the error between the measured and predicted cross-context correlation (**Fig 3d,e**). For the example HG electrode, the cross-context correlation was best-predicted by an integration period with a narrow width (68 ms) and early center (64 ms) compared with the STG electrode, which was best-predicted by a wider and more delayed integration period (375 ms width, 273 ms center). These results validate our qualitative observations and provide us with a quantitative estimate of each electrode’s integration period.

### Anatomical organization

We identified 190 electrodes with a reliable response to sound across 18 patients (test-retest correlation > 0.1; p < 10^−5^ via a permutation test across sound sequences; 128 left hemisphere; 62 right hemisphere). From these electrodes, we created a map of integration widths and centers, discarding a small fraction of electrodes (5%) where the model predictions were not highly significant (p < 10^−5^ via a phase-scrambling analysis) (**Fig 4a**). This map was created by localizing each electrode on the cortical surface, and aligning each subject’s brain to a common anatomical template. By necessity, we focus on group analyses due to the sparse, clinically-driven coverage in any given patient. Most sound-responsive electrodes were in and around the lateral sulcus and STG, as expected^11,15^.

**Fig 4.**
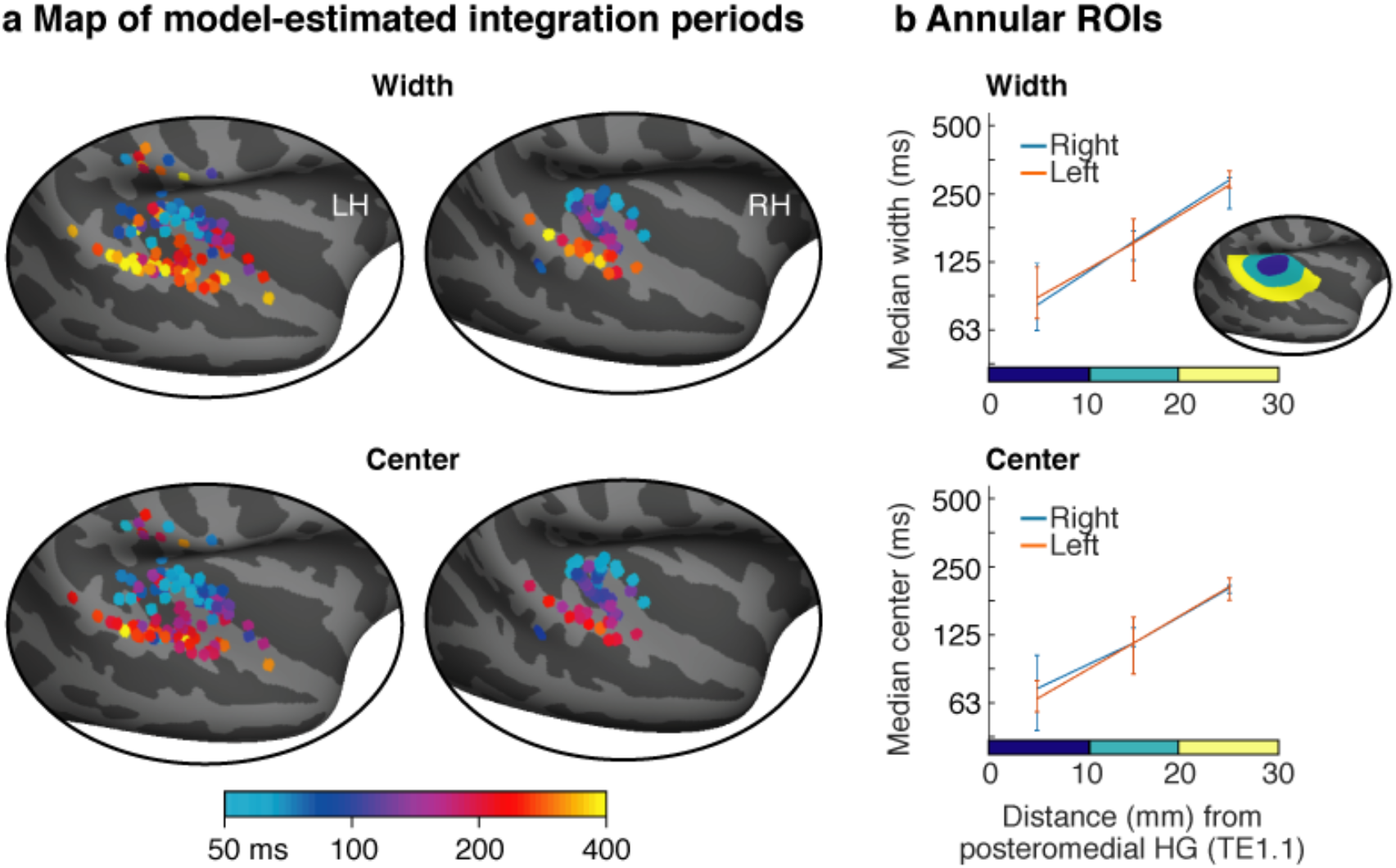
Anatomy of model-estimated integration periods. **a**, Map of integration widths (top) and centers (bottom) for all electrodes with a reliable response to sound. **b,** Electrodes were binned into ROIs based on their distance to a common anatomical landmark of primary auditory cortex (posteromedial Heschl’s gyrus, TE1.1). This figure plots the median integration width and center across the electrodes in each bin. Inset shows the ROIs for one hemisphere. Error bars plot one standard error of the bootstrapped sampling distribution across subjects.

These maps revealed a clear anatomical gradient: integration widths and centers increased substantially from primary regions near posteromedial HG to non-primary regions near STG. We quantified this trend by binning electrodes into anatomical regions-of-interest (ROIs) based on their distance to posteromedial HG (**Fig 4b**)^31^. This analysis revealed a three-fold increase in integration widths and centers from primary to non-primary regions (median integration width: 84, 152, 281 ms; median integration center: 68, 115, 203 ms; p < 0.001 via a bootstrapping analysis across subjects comparing the nearest and farthest bins). By contrast, there was no difference in integration widths or centers between the two hemispheres either when averaging across all ROIs or comparing individual ROIs (all ps > 0.74). These findings were robust across the specific sounds tested (**Fig S3**), the type of context used to assess invariance (random vs. natural; **Fig S4**), the shape of the model window (**Fig S5**), and the frequency range used to measure broadband gamma (**Fig S6**). These results demonstrate human auditory cortex integrates across time hierarchically, with substantially wider and more delayed integration periods in higher-order regions, but no difference between hemispheres.

Across all electrodes, we found that integration centers scaled approximately linearly with integration widths (**Fig S7**). In part as a consequence of this observation, we found that integration centers were relatively close to the minimum possible for a causal window even when not explicitly constrained to be causal (**Fig S7**) (integration centers were on average 46% greater than the minimum for a Gamma-distributed window). Since the integration center reflects the delay between the integration period and the response, this finding suggests that auditory cortex responds to sounds about as quickly as possible given the integration period^32^.

### Category selective responses are limited to electrodes with long integration periods

What is the consequence of hierarchical temporal integration for the analysis of important sound categories like speech and music? While prior studies have revealed non-primary neural populations that respond selectively to sound categories like speech and music^9–13^, it is unclear how these neural populations integrate information in speech and music. A priori it seemed possible that speech and music-selective responses might have diverse integration periods since sound categories like speech and music have unique structure at many timescales. For example, the median duration of speech phonemes in the popular TIMIT corpus is 64 milliseconds while syllables and words are typically hundreds of milliseconds (the median duration of multiphone syllables and multisyllable words is 197 and 479 milliseconds, respectively) (**Fig S1**). But the hierarchy revealed by our integration period maps suggests an alternative hypothesis: that category-selective responses are limited to neural responses with wide integration periods. We sought to directly test this hypothesis, and if true, determine the shortest integration periods at which category-selective responses are present.

To assess category selectivity, we ran a separate experiment in a subset of 11 patients, where we measured responses to a larger set of 119 natural sounds, drawn from 11 categories (listed in **Fig 5a**). We grouped the electrodes from these patients based on their integration width in octave intervals (shown in **Fig 5a**), pooling across both hemispheres because we had fewer electrodes (104) and because integration periods (**Fig 4**) and category-selective responses^10–12^ are similar across hemispheres. We then used several different analyses to measure the degree of category selectivity in each electrode group.

**Fig 5.**
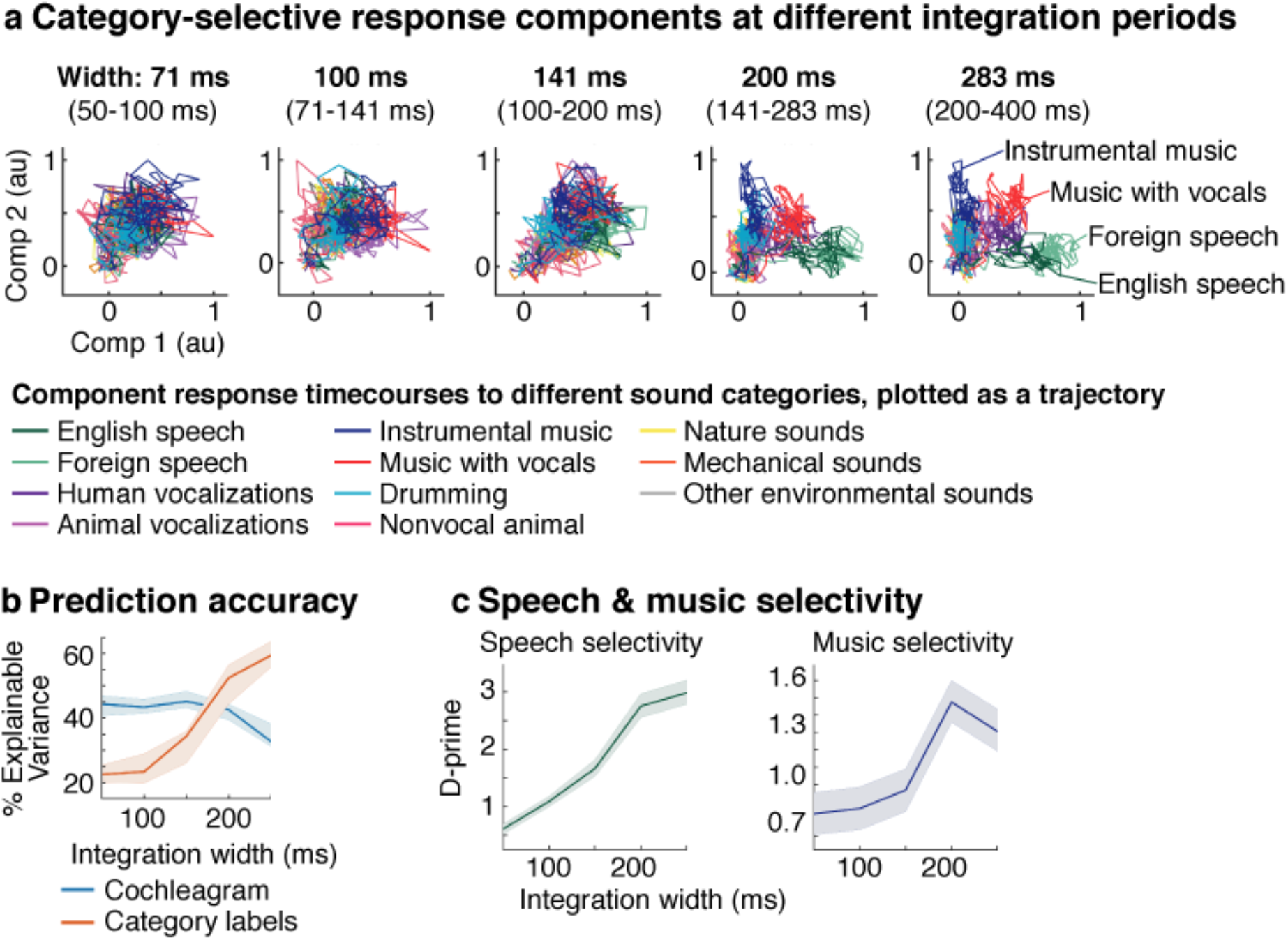
Category selectivity at different integration period widths. Responses were measured in a subset of patients to a larger collection of 119 natural sounds from 11 different sound categories, listed in panel a. Electrodes from these patients were grouped based on the width of their integration period in octave intervals. We used several different analyses to assess the degree of category-selectivity in each group. **a**, The responses from each group were projected onto the two components that exhibited the greatest category selectivity. We plot the timecourses for these two components as a 2D trajectory after averaging across the sounds from each category. Category selectivity, if present, will cause the trajectories to separate from each other. **b**, The accuracy of category labels (red line) and cochleagrams (blue line) in predicting electrode responses as a function of the integration width. This figure plots the squared correlation (noise-corrected) between the measured and predicted response for each feature set. **c**, The selectivity of each electrode group for either speech (English and foreign) or music (instrumental, vocal and drumming), measured along the component that exhibited the greatest speech or music selectivity. Selectivity was measured as the separation (noise-corrected d-prime) between responses to sounds from the target category (speech or music) compared with all other sounds. Independent sounds were used to estimate components and measure their response. Error bars plot one standard error of the bootstrapped sampling distribution.

First, we used component methods to visualize any category selectivity present in the electrodes^11^. Specifically, we projected the responses from each group onto the two components that showed the greatest category selectivity (**Fig 5a**) (estimated using cross-validated linear discriminant analysis). We plot the timecourse of these two components as a 2D trajectory after averaging across the sounds from each category. Category selectivity, if present, will cause the trajectories for different categories to separate from each other. This analysis revealed that only electrodes with wide integration periods, above ~200 milliseconds, show robust category selectivity. Similar results were obtained when analyzing the top two principal components without any optimization for category selectivity (**Fig S8**), which demonstrates that category selectivity is a prominent feature of cortical responses with wide integration periods.

To quantify these trends, we measured how accurately we could linearly predict the response of each electrode from either a cochleagram or binary category labels (**Fig 5b**). Cochleagrams are similar to spectrograms but are computed using filters designed to mimic the pseudo-logarithmic frequency resolution of cochlear filtering^30^. This analysis thus provides an estimate of the fraction of the response that can be predicted using a linear spectrotemporal receptive field^15,19,20^. The category labels indicated the membership of each sound in each category for all timepoints with sound energy above a minimum threshold. Prediction accuracies were noise-corrected using the test-retest reliability of the electrode responses, which provides an upper bound on the fraction of the response explainable by any model^20^.

We found that the prediction accuracy of the category labels more than doubled as integration widths increased, with a relatively sharp increase at ~200 milliseconds (p < 0.001 via bootstrapping across subjects). In contrast, the prediction accuracy of cochlear features decreased (p < 0.05), yielding a significant interaction between feature type and integration width (p < 0.001). This finding confirms our observation that responses become substantially more category-selective as integration periods widen.

We note the absolute prediction accuracies were modest for both the cochleagram and category labels, never exceeding more than 45% and 60% of the explainable response variance, respectively (as expected). This fact illustrates the utility of having a model-independent way of estimating integration periods, since even our best-performing models fail to explain a large fraction of the response, and the best-performing model can vary across electrodes.

The component trajectories (**Fig 5a**) suggested that selectivity for both speech and music increase as integration periods widen. To directly test this hypothesis, we separately measured the degree of speech and music selectivity at different integration widths. Selectivity was measured as the average separation (d-prime) between responses to speech or music vs. all other sounds, along the component that showed the greatest speech or music selectivity (**Fig 5c**; **Fig S9** shows the response timecourse of these components to each category). Speech sounds comprised both English and foreign speech, since we found they produced similar response trajectories, consistent with prior work showing that speech-selective responses in STG are not driven by linguistic meaning^11,12^. Non-speech sounds comprised all categories except vocal music, which produced an intermediate response along the speech-selective component, likely due to speech in the vocals (**Fig S9**). Music sounds included instrumental music, vocal music and drumming, since we have found that all three produce higher-than-average responses in music-selective regions^33^. We found that selectivity for both speech and music increased as integration periods widen (p < 0.001 via bootstrapping). This increase was more prominent for speech than music (note the different y-axes in **Fig 5c**), plausibly because speech-selective responses are more prominent particularly in the posterior/middle STG where many of our electrodes were located^11,12,15^. But for both speech and music, there was a marked increase in selectivity starting at ~200 milliseconds. This finding demonstrates that increased category selectivity is a general feature of neural responses with wide integration periods that applies to multiple sound categories.

## Discussion

Our study reveals how the human auditory cortex integrates across multiple timescales when processing natural sounds. Our findings resolve a longstanding debate by showing that auditory cortex integrates across time hierarchically, with substantially longer integration periods in non-primary regions, but no difference between hemispheres. Our results also reveal the significance of hierarchical temporal integration for the analysis of important sound categories like speech and music. In particular, we found that category-selective neural responses are restricted to electrodes with long integration periods above ~200 milliseconds. This finding suggests that short-term structure in speech and music is analyzed by general-purpose acoustic representations in primary auditory cortex and then integrated over long timescales to form category-specific representations in non-primary regions.

These findings were enabled by a novel method that makes it possible to estimate the integration period of any sensory response. Unlike prior methods, TCI makes no assumptions about the type of response being measured; it simply estimates the time window when stimuli alter the neural response. As a consequence, the method should applicable to any modality, stimulus set, or recording method. We applied TCI to intracranial recordings from epilepsy patients, using surface and depth electrodes placed throughout human auditory cortex. As a consequence, we were able to determine how anatomically and functionally distinct regions of the human brain collectively integrate across multiple timescales.

### Implications for multiscale temporal integration

Temporal integration plays a central role in most theories and models of auditory processing^2–5,7^. But it has remained unclear how the human auditory cortex integrates across multiple timescales, because there has been no general method for estimating neural integration periods.

Hemispheric models posit that the left and right hemisphere are specialized for integrating across distinct timescales^2,5^, in part to represent the distinctive temporal structure of sound categories like speech and music^6^. Some of the clearest evidence for hemispheric specialization comes from recent studies that have filtered-out temporal modulations at different rates in natural stimuli, and shown that removing fast temporal modulations has a greater impact on responses in the left auditory cortex, particularly when processing speech^6,23^. However, the time window that a neuron integrates over cannot be determined from its modulation tuning. For example, a neuron could respond selectively to fast temporal modulations over a long temporal window or to a complex structure (e.g. word) that is poorly described by its modulation content.

Another common proposal is that the auditory cortex integrates across time hierarchically^3,4,7^. Early evidence for hierarchical temporal organization came from the observation that “phase-locking” slows from the periphery to the cortex^34,35^, which implies that neurons encode temporal modulations via changes in firing rate rather than responding at particular modulation phases. But the integration period of a nonlinear response cannot be inferred from the presence or absence of phase-locking. For example, a neuron could integrate over several cycles of a modulation and then respond at the predicted peak or trough of the oscillation (phase-locking) or modulate its firing rate based on the oscillation period (rate coding).

In auditory cortex, single-unit recordings in ferrets have revealed a slight temporal broadening of linear spectrotemporal receptive fields in non-primary vs. primary auditory cortex (36 vs. 33 ms between PEG and A1)^36^. But the overall integration period cannot be inferred from these estimates since cortical responses exhibit prominent nonlinearities^20,22^, particularly in non-primary regions^31^ and particularly when responding to natural sounds^37^. In humans, several studies have revealed selectivity for naturalistic temporal structure in non-primary regions. Examples include selectivity for phonotactic structure above and beyond tuning for individual phonemes^32,38,39^, and selectivity for natural speech compared with temporally scrambled speech^12,18,25^. But again, integration periods cannot be inferred from this type of data. For example, primary regions respond similarly to intact and scrambled stimuli, even for stimuli that are scrambled at timescales well below the integration period of the neural response (e.g. 30 milliseconds)^12^. As a consequence, there has been no way to test if primary and non-primary regions differ in their integration period, and if so, by how much.

Because of these methodological limitations, there are no estimates of integration periods in human auditory cortex, making it impossible to test how auditory cortex integrates over multiple timescales. Our study thus resolves a longstanding debate by showing that multiscale temporal integration is predominantly hierarchical and not hemispheric. These results do not imply that there is no hemispheric organization for modulation tuning, since as already noted, one cannot infer the integration period of a neural response from its modulation tuning, or vice versa. Indeed, integration periods are useful specifically because they abstract away from the particular features which drive a neural response. As a consequence, integration periods can be used to compare any two brain regions, even if they respond to very different stimulus features, like primary and non-primary auditory cortex.

The hierarchical organization of temporal integration periods appears analogous to the hierarchical organization of spatial receptive fields in visual cortex^40,41^, which suggests there might be general principles that underlie this type of organization. For example, both auditory and visual recognition become increasingly challenging at large temporal and spatial scales, because the dimensionality of the input grows exponentially. Hierarchical multiscale analysis may help overcome this exponential expansion by allowing sensory systems to gradually recognize large-scale structures as combinations of smaller-scale structures (e.g. a face from nose and eyes, or a word from several phonemes) rather than attempting to recognize large-scale structures directly from the high-dimensional input^3,4,7^.

### Implications for the analysis of sound categories

Sound categories like speech and music have unique acoustic structure at many temporal scales^1,3,8,42^, from tens to hundreds of milliseconds (**Fig S1**). Prior studies have revealed non-primary neural populations selective for important sound categories like speech and music^9–13^. But very little is known about how information in speech and music is integrated in these neural populations. A prior fMRI study used a scrambling technique called “quilting” to show that speech-selective regions respond selectively to intact temporal structure up to about 500 milliseconds in duration^12^. But this study was only able to identify a single analysis timescale across all of auditory cortex, likely because scrambling is a coarse manipulation and fMRI a coarse measure of the neural response.

We were able to identify a broad range of integration periods from tens to hundreds of milliseconds, and could thus test whether category-selective responses were present at both short and long integration periods. Our results indicate that category-selective responses are only robustly present at integration periods above ~200 milliseconds, which corresponds to about the duration of a multi-phone syllable (**Fig S1**) (the duration of musical structures is less stereotyped and thus harder to assess). This finding suggests that short-term structures (e.g. phonemes) are analyzed by general-purpose acoustic representations in primary auditory cortex (e.g. spectrotemporal receptive fields)^16^, and then integrated over long timescales to form category-specific representations in non-primary regions. This finding does not imply that speech-selective regions are insensitive to short-term structure such as phonemes, but rather that speech-selective responses respond to larger-scale patterns, such as phoneme sequences, consistent with recent work on phonotactics^32,38,39^.

We found that both speech and music selectivity increased for integration periods greater than ~200 milliseconds (**Fig 5c**), which is perhaps surprising given that speech and music have distinctive temporal structure^2,43^. This result might be explained by the fact that speech and music-selective responses emerge at a similar point in the cortical hierarchy^10,11^, just beyond primary auditory cortex. If integration periods are predominantly organized hierarchically, as suggested by our data, regions with a similar hierarchical position might exhibit similar integration periods even if they respond to different stimuli.

### Limitations and extensions

As with any method, our results could depend upon the stimuli tested. We tested a diverse set of natural sounds with goal of characterizing responses throughout auditory cortex using ecologically relevant stimuli. Because time is short when working with surgical patients, we could only able to test a small number of sounds, but found that our key findings were robust to the sounds tested (**Fig S3**). Nonetheless, it will be important in future work to test whether and how integration periods change for different stimulus classes.

Another key question is whether temporal integration periods reflect a fixed property of the cortical hierarchy or whether they are shaped by attention and behavioral demands. In our study, we did not give subjects a formal task because our goal was to measure integration periods during natural listening without any particular goal or attentional focus. Future work could explore how behavioral demands shape temporal integration periods by measuring integration periods in the presence or absence of focused attention to a short-duration (e.g. phoneme) or long-duration (e.g. word) target^44^.

Temporal integration periods indicate the time window in the stimulus that a neural response is sensitive to, which is distinct from the time window in the neural response that best encodes a property of the stimulus, sometimes referred to as the “encoding window”^17,45–47^. For example, the encoding window for a slow temporal modulation could be quite long, even if the neural integration period is quite short. Encoding windows thus reflect a complex mixture of the neural integration period and the temporal properties of the stimulus being encoded.

A given neural response might effectively have multiple integration periods. For example, neural responses are known to adapt their response to repeated sounds on the timescale of seconds^48^ to minutes^49^ and even hours^50^, suggesting a form of long-term memory^51^. TCI measures the integration period of responses that are reliable across repetitions, and as a consequence, TCI will be insensitive to response characteristics that change across repeated presentations. Future work could try and identify multiple integration periods within the same response by manipulating the type of context which surrounds a segment. Here, we examined two distinct types of contexts and found similar results (**Fig S4**), suggesting that hierarchical temporal integration is a robust property of human auditory cortex.

Our study focused on characterizing responses within auditory cortex using a relatively short range of segment durations (31 milliseconds to 2 seconds) and a diverse set of natural sounds, including many non-speech and non-music sounds. Future work could characterize integration periods outside of auditory cortex by using a longer range of segment durations and focusing on just speech or music, which exhibit longer-term temporal structure^8,18,42^. For example, a recent fMRI study provided evidence for multi-second integration periods in regions outside of auditory cortex by examining the delay needed for responses to speech to become context invariant^52^. This study was not able to estimate temporal integration periods within auditory cortex because the timescale of the fMRI response is an order of magnitude slower than auditory cortical integration periods. Here, we took advantage of the rare opportunity to record intracranial responses from the human brain, which is the only human neuroscience method with the spatiotemporal resolution to estimate integration periods in auditory cortex.

## Methods

### Participants & data collection

Data were collected from 23 patients undergoing treatment for intractable epilepsy at the NYU Langone Hospital (14 patients) and the Columbia University Medical Center (9 patients). One patient was excluded because they had a large portion of the left temporal lobe resected in a prior surgery. Of the remaining 22 subjects, 18 had sound-responsive electrodes (see *Electrode selection*). Electrodes were implanted to localize epileptogenic zones and delineate these zones from eloquent cortical areas before brain resection. NYU patients were implanted with subdural grids, strips and depth electrodes depending on the clinical needs of the patient. CUMC patients were implanted with stereotactic depth electrodes. All subjects gave informed written consent to participate in the study, which was approved by the Institutional Review Boards of CUMC and NYU.

### Stimuli for TCI paradigm

Segments were excerpted from 10 natural sound recordings, each two seconds in duration (**Table S1**). Shorter segments were created by subdividing the longer segments. Each natural sound was RMS-normalized before segmentation.

We tested seven segment durations (31.25, 62.5, 125, 250, 500, 1000, and 2000 ms). We presented the segments of a given duration in two pseudorandom orders, yielding 14 sequences (7 durations × 2 orders), each 20 seconds in duration. The only constraint was that a given segment had to be preceded by a different segment in the two orders. When we designed the stimuli, we thought that integration periods might be influenced by transients at the start of a sequence, so we designed the sequences such that the first 2 seconds and last 18 seconds of each sequence contained distinct and non-overlapping segments so that we could separately analyze the just last 18 seconds. In practice, integration periods were similar when analyzing the first 18 seconds vs. the entire 20-second sequence. Segments were concatenated using cross-fading to avoid click artifacts (31.25 ms raised cosine window; cross-fading was accounted for in our integration period model). Each stimulus was repeated several times (4 repetitions for most subjects; 8 repetitions for 2 subjects; 6 and 3 repetitions for two other subjects). Stimuli will be made available upon publication.

### Natural sounds

In a subset of 11 patients, we measured responses to a diverse set of 119 natural sounds from 11 categories, similar to those from our prior studies characterizing auditory cortex^11^ (there were at least 7 exemplars per category). The sound categories are listed in **Figure 5a**. Most sounds (108) were 4 seconds in duration. The remaining 11 sounds were longer excerpts of English speech (28-70 seconds) that were included to characterize responses to speech for a separate study. Here, we just used responses to the first 4 seconds of these stimuli to make them comparable to the others. The longer excerpts were presented either at the beginning (6 patients) or end of the experiment (5 patients). The non-English speech stimuli were drawn from 10 languages: German, French, Italian, Spanish, Russian, Hindi, Chinese, Swahili, Arabic, Japanese. We classified these stimuli as “foreign speech” since nearly all were unfamiliar to our subjects, though occasionally a patient had some familiarity with one language. Twelve sounds were repeated four times to make it possible to measure response reliability and noise-correct our measures. All other stimuli were presented once. All sounds were RMS normalized.

As with the main experiment, subjects did not have a formal task but the experiment was periodically paused and subjects were asked a simple question to encourage them to listen to the sounds. For the 4-second sounds, subjects were asked to identify/describe the last sound they heard. For the longer English speech excerpts, subjects were asked to repeat the last phrase they heard.

### Preprocessing

Electrode responses were common-average referenced to the grand mean across electrodes from each subject. We excluded noisy electrodes from the common-average reference by detecting anomalies in the 60 Hz power band (measured using an IIR resonance filter with a 3dB down bandwidth of 0.6 Hz). Specifically, we excluded electrodes whose 60 Hz power exceeded 5 standard deviations of the median across electrodes. Because the standard deviation is itself sensitive to outliers, we estimated the standard deviation using the central 20% of samples, which are unlikely to be influenced by outliers. Specifically, we divided the range of the central 20% of samples by that which would be expected from a Gaussian of unit variance. After common-average referencing, we used a notch filter to remove harmonics & fractional multiples of the 60 Hz noise (60, 90, 120, 180; using an IIR notch filter with a 3dB down bandwidth of 1 Hz; the filter was applied forward and backward).

We computed broadband gamma power by measuring the envelope of the preprocessed signal filtered between 70 and 140 Hz (implemented using a 6^th^ order Butterworth filter with 3dB down cutoffs of 70 and 140 Hz; the filter was applied forward and backward). Results were similar using other frequency ranges (**Fig S6**). The envelope was measured as the absolute value of the analytic signal after bandpassing. We then downsampled the envelopes to 100 Hz (the original sampling rate was either 512, 1000, 1024, or 2048 Hz, depending on the subject). We used simulations to ensure that this procedure would allow us to accurately recover the integration period of broadband power fluctuations (see *Simulations* below).

Occasionally, we observed visually obvious artifacts in the broadband gamma power for a small number of timepoints. To detect such artifacts, we computed the 90^th^ percentile of each electrode’s response distribution across all timepoints. We classified a timepoint as an outlier if it exceeded 5 times the 90^th^ percentile value for each electrode. We found this value to be relatively conservative in that only a small number of timepoints were excluded (on average, 0.04% of timepoints were excluded across all sound-responsive voxels). Because there were only a small number of outlier timepoints, we replaced the outlier values with interpolated values from nearby non-outlier timepoints.

As is standard, we time-locked the iEEG recordings to the stimuli by either cross-correlating the audio with a recording of the audio collected synchronously with the iEEG data or by detecting a series of pulses at the start of each stimulus that were recorded synchronously with the iEEG data. We used the stereo jack on the experimental laptop to either send two copies of the audio or to send audio and pulses on separate channels. The audio on one channel was used to play sounds to subjects, and the audio/pulses on the other were sent to the recording rig. Sounds were played through either a Bose Soundlink Mini II speaker (at CUMC) or an Anker Soundcore speaker (at NYU). Responses were converted to units of percent signal change relative to silence by subtracting and then dividing the response of each electrode by the average response during the 500 ms before each stimulus.

### Electrode selection

We selected electrodes with a reliable broadband gamma response to the sound set. Specifically, we measured the test-retest correlation of each electrodes response across all stimuli (using odd vs. even repetitions). We selected electrodes with a test-retest Pearson correlation of at least 0.1, which we found to be sufficient to reliably estimate integration periods in simulations (described below). We ensured that this correlation value was significant using a permutation test, where we randomized the mapping between stimuli across repeated presentations and recomputed the correlation (using 1000 permutations). We used a Gaussian fit to the distribution of permuted correlation coefficients to compute small p-values^53^. Only electrodes with a highly significant correlation relative to the null were kept (p < 10^−5^). We used a low p-value threshold, because we have found that any electrode with a borderline-significant test-retest response across an entire experiment is very noisy. We identified 190 electrodes out of 2847 total that showed a reliable response to natural sounds based on these criteria.

### Denoising

Responses in non-primary regions were less reliable on average then responses in primary regions (the median test-retest correlation for each annular ROIs in **Fig 4b** was 0.32, 0.23, 0.17). We ensured that differences in reliability could not explain our results in two ways: (1) we ensured that our model-estimated integration periods were unbiased by low data reliability (see *Model-estimated integration periods* and *Simulations* below) (2) we repeated our analyses using a denoising procedure that substantially increased the reliability of the electrode responses to a level well-above the point at which reliability might affect our integration period estimates. Our denoising procedure was motivated by the observation that iEEG responses are relatively low-dimensional and as a consequence much of the stimulus-driven response variation is shared across subjects^54^, in contrast with the noise which differs from subject to subject. We thus projected the electrode responses from one subject onto the responses from all other subjects (using regression), which has the effect of throwing out response variation that is not present in at least two subjects. We have found this procedure to be useful when there are a relatively large number of subjects with responses from a restricted region of the brain like auditory cortex, as was the case in our study. To examine the effect of denoising, we measured split-half reliability before and after denoising (**Fig S10a**). When we denoised both splits of data, the median correlation increased four-fold from 0.21 to 0.8 (**Fig S10a**, purple dots). We also found that the reliability improved when only one split was denoised, which indicates that the analysis discarded more noise than signal (reliability improved for 93% of electrodes) (**Fig S10a**, blue dots). Since our results were similar using original and denoised data (compare **Figs 2b&4** with **Figs S10b&c**), we conclude that our findings cannot be explained by differences in data reliability.

### Electrode localization

Following standard practice, we localized electrodes as bright spots on a post-operative computer tomography (CT) image or dark spots on a magnetic resonance image (MRI), depending on whichever was available in a given patient. The post-op CT or MRI was aligned to a high-resolution, pre-operative magnetic resonance image (MRI) that was undistorted by electrodes. Each electrode was then projected onto the cortical surface computed by Freesurfer from the pre-op MRI scan, excluding electrodes that were greater than 10 mm from the surface. This projection is error prone because faraway points on the cortical surface can be nearby in space due to cortical folding. To minimize gross errors, we preferentially localized sound-responsive electrodes to regions where sound-driven responses are likely to occur^54^. Specifically, we calculated the likelihood of observing a significant response to sound using a recently collected fMRI dataset, where responses were measured to a large set of natural sounds across 20 subjects with whole-brain coverage^33^ (p < 10^−5^, measured using a permutation test). We treated this map as a prior and multiplied it by a likelihood map, computed separately for each electrode based on the distance of that electrode to each point on the cortical surface (using a 10 mm FWHM Gaussian error distribution). We then assigned each electrode to the point on the cortical surface where the product of the prior and likelihood was greatest (which can be thought of as the maximum posterior probability solution). We smoothed the prior probability map (10 mm FWHM kernel) so that it would only affect the localization of electrodes at a coarse level, and not bias the location of electrodes locally, and we set the minimum prior probability to be 0.05 to ensure every point had non-zero prior probability. We plot the prior map and its effect on localization in **Fig S11**.

### Cross-context correlation

We now review our key analysis for estimating context invariance. MATLAB code implementing these analyses will be made available upon publication. For each electrode and segment duration, we compiled the responses surrounding all segments into a matrix, aligned to segment onset (the segment-aligned response matrix or SIR matrix) (**Fig 2a**). We calculated a separate SIR matrix for each context, such that corresponding rows contained the response timecourse to the same segment from different contexts. To detect if there was a lag where the response was the same across contexts, we correlated corresponding columns across SIR matrices from different contexts (the cross-context correlation). We compared the cross-context correlation with the correlation when the context was identical using repeated presentations of the same sequence (the noise ceiling).

The noise ceiling exhibited reliable variation across lags, which is evident from the fact that the cross-context correlation remained yoked to the noise ceiling when the segment duration was long relative to the integration period (evident for example in the HG electrode’s data for 250 and 500 ms in **Fig 2b**). This variation is expected since the sounds that happen to fall within the integration period will vary with lag, and the noise ceiling will depend upon how strongly the electrode responds to the sounds within the integration period. It is also evident that the variation in the noise ceiling is periodic. This periodicity is an inevitable consequence of correlating across a fixed set of segments. To see this, consider the fact that the onset of one segment is also the offset of the preceding segment. Since we are correlating across segments for a given lag, the values being used to compute the correlation are nearly identical at the start and end of a segment (the only difference occurs for the first and last segment of the entire sequence). The same logic applies to all lags that are separated by a period equal to the segment duration.

Because the shorter segments were subsets of the longer segments, we could consider two types of context: (1) random context, where a segment is flanked by random other segments (2) natural context, where a segment is a part of a longer segment and thus surrounded by its natural context. Since the two contexts being compared must differ, one of the contexts always has to be random, but the other context can be random or natural. In practice, we found similar results when comparing random-only contexts and when comparing random and natural contexts (**Fig S4**). This fact is practically useful since it greatly increases the number of comparisons that can be made. For example, each 31 millisecond segment had 2 random contexts (one per sequence) and 12 natural contexts (2 sequences × 6 longer segment durations). The two random contexts can be compared with each other as well as with the other 12 natural contexts. For our main analyses, we averaged the cross-context correlation across all of these comparisons for maximal statistical power.

We note that the cross-context correlation will typically be more reliable for shorter segment durations since there are more segments with which to compute the correlation. We consider this property useful since for electrodes with shorter integration periods there will be a smaller number of lags at the shorter segment durations that effectively determine the integration period, and thus it is useful if these lags are more reliable. Conversely, electrodes with longer integration periods exhibit a more gradual build-up of the cross-context correlation at the longer segment durations, and our model enables us to pool across all of these lags to arrive at a robust estimate of the integration period.

### Model-estimated integration periods

We modeled temporal integration periods using a Gamma-distributed window (*h*) that we scaled and shifted in time:

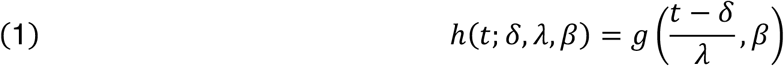

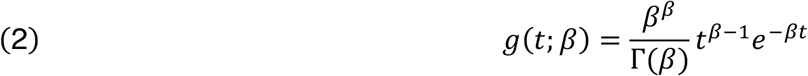

The shape is determined by *β* and varies from more exponential to more bell-shaped (**Fig S5**). The integration width and center do not correspond directly to any of the three parameters (*δ*, *λ*, *β*), mainly because the scale parameter (*λ*) alters both the center and width. The integration width was defined as the smallest interval that contained 75% of the window’s mass, and the integration center was defined as the window’s median. Both parameters were calculated numerically from the cumulative distribution function of the window.

For a given integration window, we predicted the cross-context correlation at each lag and segment duration by measuring how much the integration period overlaps the shared central segment (*w*) vs. the N surrounding context segments (*β*_*n*_) (see **Fig 3b**):

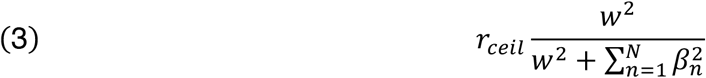

where *r*_*ceil*_ is the measured noise ceiling, and the ratio on the right is the predicted correlation in the absence of noise. The predicted cross-context correlation varies with the segment duration and lag because the overlap varies with the segment duration and lag. When the integration period only overlaps the shared segment (*w* = 1, Σ*β*_*n*_ = 0), the model predicts a correlation of 1 in the absence of noise, and when the integration period only overlaps the surrounding context segments (*w* = 0, Σ*β*_*n*_ = 1), the model predicts a correlation of 0. In between these two extremes, the predicted cross-context correlation equals the fraction of the response driven by the shared segment, with the response variance for each segment given by the squared overlap with the integration period. A formal derivation of this equation is given at the end of the Methods (see *Deriving a prediction for the cross-context correlation*). For a given segment duration, the overlap with each segment was computed by convolving the model integration period with a boxcar function whose width is equal to the segment duration (with edges tapered to account for cross-fading).

We varied the width, center and shape of the model integration period and selected the window with the smallest prediction error. Since the cross-context correlation is more reliable for shorter segment durations due to the greater number of segments, we weighted the error by the number of segments used to compute the correlation before averaging across segment durations. Integration widths varied between 31.25 and 1 second (using 100 logarithmically spaced steps). Integration centers varied from the minimum possible given for a causal window up to 500 milliseconds beyond the minimum in 10 millisecond steps. We tested five window shapes (*β* = 1,2,3,4,5).

We found in simulations that there was an upward bias in the estimated integration widths for noisy data when using the mean squared error (see *Simulations* below). We checked that this bias did not affect our results in two ways. First, we repeated our analyses using denoised data, whose reliability was well above the point at which the bias has any effect (**Fig S10**). Second, we derived a bias-corrected metric, which substantially reduced the bias in simulations (see *Bias correction* below). We used this bias-corrected metric for all of our analyses, but found that the results were very similar with and without bias correction, indicating that our data were sufficiently reliable to avoid any substantial bias even without denoising (compare **Fig 4** which shows results with correction with **Fig S12** which shows results without correction).

We assessed the significance of our model predictions by creating a null distribution using phase-scrambled model predictions. Phase scrambling exactly preserves the mean, variance and autocorrelation of the predictions but alters the locations of the peaks and valleys. Phase scrambling was implemented by shuffling the phases of different frequency components without altering their amplitude and then reconstructing the signal (using the FFT/iFFT). After phase-scrambling, we remeasured the error between the predicted and measured cross-context correlation, and selected the model with the smallest error (as was done for the unscrambled predictions). We repeated this procedure 100 times to build up a null distribution, and used this null distribution to calculate a p-value for the actual error based on unscrambled predictions (again fitting the null distribution with a Gaussian to calculate small p-values). For 95% of sound-responsive electrodes (181 of 190), the model’s predictions were highly significant (p < 10^−5^).

### Simulations

We tested our complete analysis pipeline using simulated data. Specifically, we modulated a broadband carrier (Gaussian noise filtered between 70 and 140 Hz) with the waveform amplitude of the stimuli from our TCI paradigm, integrated within a Gamma-distributed integration period. We added Gaussian noise to manipulate the test-retest reliability of these responses and thus determine the minimum reliability needed to accurately infer integration periods. We simulated four different responses to the same stimuli using independent samples of the broadband carrier and additive noise (for most subjects we had four repetitions), and iteratively increased or decreased the noise level to achieve a desired split-half correlation (within a tolerance of 0.001).

We then applied our complete analysis pipeline to these simulated responses. Our goal was to assess if we could accurately infer the true integration width from the simulated data. We thus varied the integration width of the simulated response (from 31 ms to 500 ms in octave steps), using a fixed shape (*β* = 3) and center (set to the minimum value for a casual window). However, we did not assume that the shape or center were known, and thus varied the shape and center along with the width when inferring the best-fit integration period, as was done for the iEEG analyses.

We found that the estimated integration widths were close to the true widths when the split-half reliability was at least 0.1 (**Fig S13**), which was the reliability cutoff used to select sound-responsive electrodes. When the test-retest reliability was low, there was an upwards bias in the estimated widths when using the mean squarer error (**Fig S13**, top panel), which was substantially reduced using our bias-corrected metric (**Fig S13**, bottom panel; see *Bias correction* below).

### Anatomical ROI analyses

We grouped electrodes into regions-of-interest (ROI) based on their anatomical distance to posteromedial Heschl’s gyrus (TE1.1)^55^ (**Fig 4b**), which is a common anatomical landmark for primary auditory cortex^31,56^. Distance was measured on the flattened 2D representation of the cortical surface as computed by Freesurfer. Electrodes were grouped into three 10 millimeter bins (0-10, 10-20, and 20-30 mm), and we measured the median integration width and center across the electrodes in each bin, separately for each of the two hemispheres.

Error bars and significance were computed by bootstrapping across subjects. Specifically, we resampled subjects with replacement 10,000 times and recomputed the median integration width or center within each ROI using the resampled dataset. A small fraction of these samples (1.2%) were discarded because the resampled dataset did not contain any electrodes in one of the six ROIs (3 distances × 2 hemispheres). To assess whether two ROIs significantly differed (e.g. nearest vs. farthest, left vs. right), we counted the fraction of samples where the resampled values were consistently higher or lower in one of the two ROIs (whichever fraction was lower), subtracted this fraction from 1, and multiplied by 2 to arrive at a two-sided p-value.

### Category-selective components at different temporal integration periods

To investigate selectivity for categories, we used responses to the larger set of 119 natural sounds that were tested in a subset of 11 patients. There were 104 electrodes from these 11 subjects that passed the inclusion criteria described above (out of 181 total). We grouped these electrodes based on the width of their integration period in octave intervals, spaced a half-octave apart (intervals shown in **Fig 5a**). Each group had between 22 and 43 electrodes. We then used several different analyses to investigate the degree of category selectivity in each group.

We used a combination of component methods (**Fig 5a,c**) and individual-electrode analyses (**Fig 5b**) to assess category selectivity. Component methods are commonly used to summarize responses from a population of electrodes or neurons^57^. And we have previously shown that component methods can better isolate selectivity for categories compared with analyzing individual iEEG electrodes^54^ or individual fMRI voxels^11^. To visualize the dominant structure at each integration period, we projected the responses onto the top two principle components (PCs) from each group of electrodes (**Fig S8**). If these components exhibit selectivity for categories then the average component response to different categories should appear segregated when plotted as a trajectory. Because the first two PCs might obscure category selectivity present at higher PCs, we repeated the analysis using the two components that best separated the categories, estimated using linear discriminant analysis (LDA)^58^ (**Fig 5a**). LDA was applied to timepoints between 250 milliseconds and 4 seconds after stimulus onset to account for response delays. To avoid statistical circularity, we used half the sounds to infer components, and the other half to measure their response. And to prevent the analysis from targeting extremely low-variance components, we applied LDA to the top five PCs from each electrode group.

PCs were computed using responses from the TCI experiment, where we had responses from a larger number of electrodes and subjects (181 electrodes from 18 subjects). We then estimated the response of these same PCs to the larger set of 119 natural sounds using the subset of electrodes with responses in both experiments (104 electrodes from 11 subjects). Since each PC is just a weighted sum of the electrode responses, we simply multiplied the responses to the 119 natural sounds by the reconstruction weights inferred from the TCI experiment. Since only a subset of electrodes were tested in both experiments, we inferred the reconstruction weights using just the electrodes tested in both experiments, by finding the linear combination of these electrodes that best approximated each PC.

### Feature predictions

As a complement to the component analyses, we measured the degree to which individual electrode responses could be predicted from category labels (**Fig 5b**). We binned the results based on integration width of the electrode, using the same octave-spaced intervals. And we compared the prediction accuracies for the category labels with those from a cochleagram representation of sound.

Cochleagrams were calculated using a cosine filterbank with bandwidths designed to mimic cochlear tuning^31^ (29 filters between 50 Hz and 20 kHz, 2x overcomplete). The envelopes from the output of each filter were compressed to mimic cochlear amplification (0.3 power). The frequency axis was resampled to a resolution of 12 cycles per octave and the time axis was resampled to 100 Hz (the sampling rate used for all of our analyses).

For each category label, we created a binary timecourse with 1s for all timepoints/sounds from that category, and 0s for all other timepoints. We only labeled timepoints with a 1 if they had sound energy that exceeded a minimum threshold. The sound energy at each moment in time was calculated by averaging the cochleagram across frequency, and the minimum threshold was set to one fifth the mean energy across all timepoints and sounds.

We predicted electrode responses between 500 milliseconds pre-stimulus onset to 4 seconds post-stimulus onset. We used ridge regression to learn a linear mapping between these features and the response. We included five delayed copies of each regressor, with the delays selected to span the integration period of the electrode (from the bottom fifth to the top fifth quintile). Regression weights were fit using the 107 sounds that were presented once, and we evaluated the fits using the 12 test sounds that were repeated four times each, making it possible to compute a noise-corrected measure of prediction accuracy^59,60^:

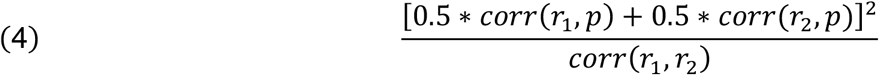

where *r*_1_ and *r*_2_ are two independent measures of the response (computed using odd and even repetitions) and *p* is the prediction computed from the training data. We used cross-validation within the training set to choose the regularization coefficient (testing a wide range of values from 2^−100^ to 2^100^ in octave steps).

Significance and error bars were calculated using bootstrapping across subjects (the same procedure described above in *Anatomical ROI analyses*). To test whether the prediction accuracy increased or decreased as a function of the integration width, we measured the slope between the noise-corrected prediction accuracy and the integration width (on a logarithmic scale). We then tested whether the bootstrapped slopes for the category and cochlear predictions differed significantly from zero and from each other.

### Speech and music selectivity

We separately quantified the degree of speech and music selectivity at each integration width (**Fig 5c**). Selectivity was quantified as the degree of separation between speech/music and all other sounds, along the components that showed the greatest selectivity for speech/music. Both English and foreign speech sounds were grouped as speech, since they yielded similar responses, consistent with prior results^11,12^. Vocal music was excluded from the speech selectivity analysis since it produced an intermediate responses along the speech-selective component (**Fig S9**), as expected since vocals contain speech^11^. Instrumental music, vocal music and drumming were grouped as music, since they produce above average responses in music-selective brain regions^33^.

The weights for each component were learned by regressing the electrode responses from each group against a binary category vector with 1s for all timepoints/sounds from the target category (e.g. speech) and 0s for all other timepoints/sounds. To avoid over-fitting to low-variance signals, we again applied our analysis to the top 5 PCs from each group. We used independent sounds to estimate components and measure their response (5-fold cross-validation, each fold had a similar number of sounds per category). To account for the response delay we only used responses between 250 milliseconds and 4 seconds post-stimulus onset.

**Figure S9** plots the average component response timecourse to each sound category (averaged across test folds). **Figure 5c** plots the average separation, measured as the d-prime between responses to sounds from the target and non-target category across all timepoints (between 250 milliseconds and 4 seconds post-stimulus onset). D-prime is a standard measure of the separation between two responses, and is defined as the difference in the mean response divided by square root of the average variances:

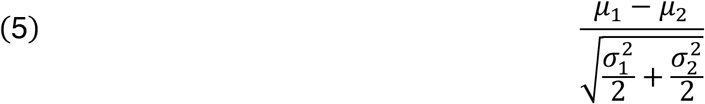

The average within-category variance in the denominator of equation 5 is a sum of the stimulus-driven response variance, which is repeatable across measurements, and the noise variance, which is not (the means are unbiased by noise assuming the noise is zero mean). We therefore noise-corrected our d-prime measure by subtracting off an estimate of the noise variance from the measured within-category variance. We estimated the noise variance as half of the error variance to repeated presentations of the same stimulus (using the 12 sounds repeated 4 times)^31^. We used half of the error variance, since the error reflects the difference between two independent measurements of the same signal, and the total variance of two independent signals that are subtracted or summed is additive.

Significance and error bars were computed via bootstrapping across sounds. Unlike other analyses, we did not bootstrap across subjects because doing so would have been inappropriate for this particular analysis. Specifically, each component was computed by regressing the electrode responses from all subjects against the target category vector, and thus in the language of regression, electrodes/subjects are features and timepoints/sounds are observations. While bootstrapping across observations (timepoints/sounds) is standard^61^, bootstrapping across features (electrodes/subjects) is inappropriate, because repeating features does not change the least-squares solution.

We bootstrapped across sounds by sampling sounds with replacement, separately for the target and non-target category (e.g. speech and non-speech sounds). We also resampled sounds from the test set used to calculate the noise variance. We then recalculated the noise-corrected d-prime using the component timecourses for the resampled sounds (repeating timecourses for sounds sampled more than once). To test if selectivity increased with the integration width, we measured if the bootstrapped slope between selectivity and the integration width was significantly greater than 0.

### Deriving a prediction for the cross-context correlation

We now derive the equation used to predict the cross-context correlation from a model integration period (equation 3). The cross-context correlation is computed across segments for a fixed lag and segment duration by correlating corresponding columns of SIR matrices from different contexts (**Fig 2a**). Consider two pairs of cells (*e*_*s,A*_, *e*_*s,B*_) from these SIR matrices, representing the response to a single segment (*s*) in two different contexts (*A*, *B*) for a fixed lag and segment duration (we do not indicate the lag and segment duration to simplify notation). To reason about how the shared and context segments might relate to the cross-context correlation at each moment in time, we assume that the response reflects the sum of the responses to each segment weighted by the degree of overlap with the integration period (**Fig 3b**):

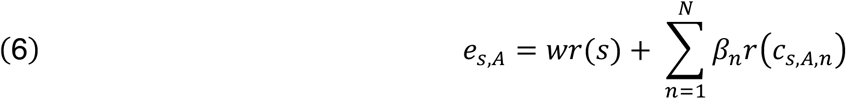

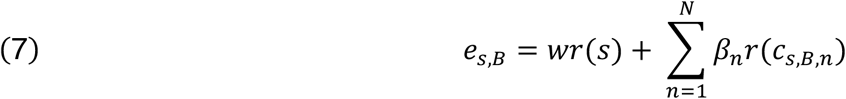

where *r*(*s*) reflects the response to the shared central segment, *r*(*C*_*s,A,n*_) and *r*(*C*_*s,B,n*_) reflect the response to the n-th surrounding segment in each of the two contexts (i.e. the segment right before and right after, two before and two after, etc.), and *w* and *β*_*n*_ reflect the degree of overlap with the shared and surrounding segments, respectively (illustrated in **Fig 3b**).

Below we write down the expectation of the cross-context correlation in the absence of noise, substitute equations 6 & 7, and simplify. Moving from line 8 to line 9 takes advantage of the fact that contexts A and B are no different in structure and so their expected variance is the same. Moving from line 10 to line 11, we have taken advantage of the fact that surrounding context segments are random, and thus all cross products that involve the context segments are zero in expectation, canceling out all of the terms except those noted in equation 11. Finally, in moving from equation 11 to 12, we take advantage of the fact that there is nothing special about the segments that make up the shared central segments compared with the surrounding context segments, and their expected variance is therefore equal and cancels between the numerator and denominator.

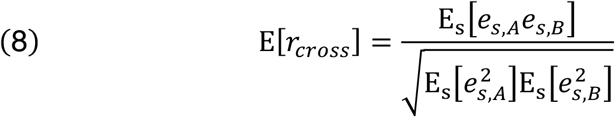

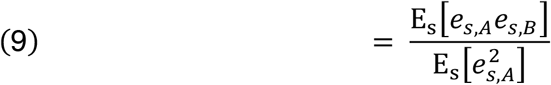

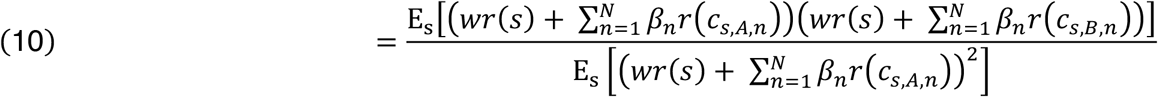

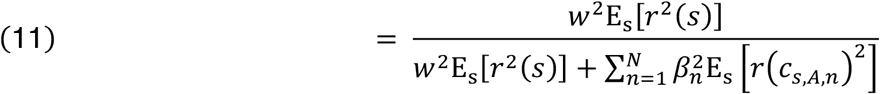

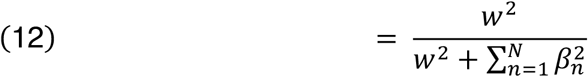

We multiplied equation 12 by the noise ceiling to arrive at our prediction of the cross context correlation (equation 3).

### Bias correction

Here, we derive the correction procedure used to minimize the bias when evaluating model predictions via the squared error.

Before beginning, we highlight a potentially confusing, but necessary distinction between noisy measures and noisy data. As we show below, the bias is caused by the fact that our correlation measures are noisy in the sense that they will not be the same across repetitions of the experiment. The bias is *not* directly caused by the fact that the data is noisy, since if there are enough segments the correlation measures will be reliable even if the data are noisy, which is what matters since we explicitly measure and account for the noise ceiling. To avoid confusion, we use the superscript (*n*) to indicate noisy measures, (*t*) to indicate the true value of a noisy measure (i.e. in the limit of infinite segments), and (*p*) to indicate a “pure” measure computed from noise-free data.

Consider the error between the measured 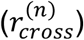 and model-predicted 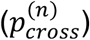 cross-context correlation for a single lag and segment duration (the model prediction is noisy because of multiplication with the noise ceiling which is measured from data):

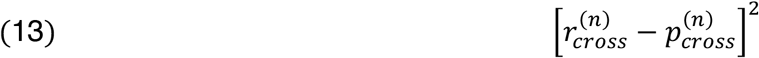

Our final cost function averaged these pointwise squared errors across all lags and segment durations weighted by the number of segments used to compute each correlation (which was greater for shorter segment durations). Here, we analyze each lag and segment duration separately, and thus ignore the influence of the weights which is simply a multiplicative factor that can be applied at the end after bias correction.

Our analysis proceeds by writing the measured 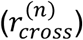 and predicted 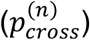 cross-context correlation in terms of their underlying true and pure measures (equations 14 to 17). We then substituting these definitions into the expectation of the squared error and simplify (equations 18 to 21), which yields insight into the cause of the bias.

The cross-context correlation 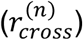 is the sum of the true cross-context correlation plus error:

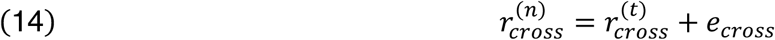

And the true cross-context correlation is the product of the pure/noise-free cross-context correlation 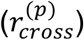 with the true noise ceiling 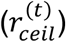:

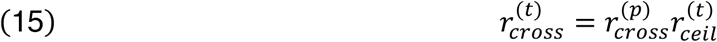

The predicted cross-context correlation is the product of the noise-free prediction 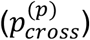 times the measured noise ceiling 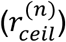:

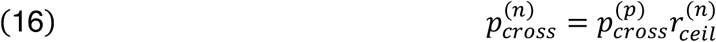

And the measured noise ceiling is the sum of the true noise ceiling 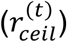 plus error 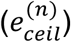 :

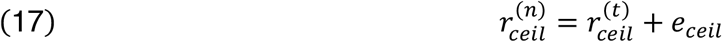

Below we substitute the above equations into the expectation for the squared error and simplify. Only the error terms (*e*_*cross*_ and *e*_*ceil*_) are random variables, and thus in equation 20, we have moved all of the other terms out of the expectation. In moving from equations 20 to 21, we make the assumption / approximation that the errors are uncorrelated and zero mean, which causes all but three terms to dropout in equation 21. This approximation, while possibly imperfect, substantially simplifies the expectation and makes it possible to derive a simple bias correction procedure, as described next.

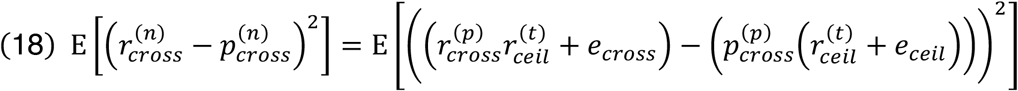

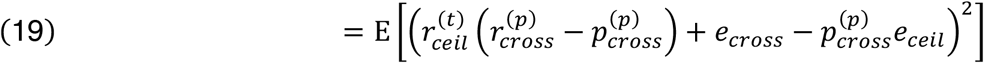

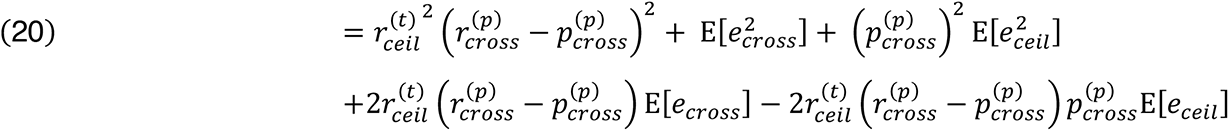

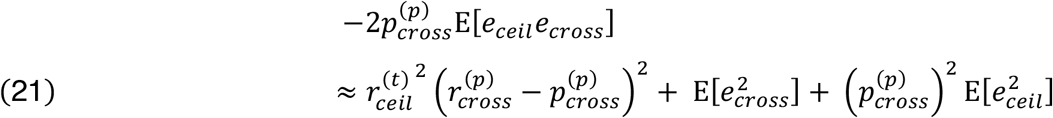

The first term in equation 21 is what we would hope to measure: a factor which is proportional to the squared error between the pure cross-context correlation computed from noise-free data 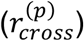 and the model’s prediction of the pure cross-context correlation 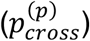. The second term does not depend upon the model’s prediction and thus can be viewed as a constant from the standpoint of analyzing model bias. The third term is potentially problematic, since it biases the error upwards based on the squared magnitude of the predictions, with the magnitude of the bias determined by the magnitude of the errors in the noise ceiling. This term results in an upward bias in the estimated integration width, because narrower integration periods have larger squared magnitudes on average. This bias is only present when there is substantial error in the noise ceiling, which explains why we only observed the bias for data with low reliability (**Fig S13**, top panel).

We can correct for this bias by subtracting a factor whose expectation is equal to the problematic third term in equation 21. All we need is a sample of the error in the noise ceiling, which our procedure naturally provides since we measure the noise ceiling separately for segments from each of the two contexts and then average these two estimates. Thus, we can get a sample of the error by subtracting our two samples of the correlation ceiling and dividing by 2 (averaging is equivalent to summing and dividing by 2 and the noise power of summed and subtracted signals is equal). We then take this sample of the error multiply it by our model prediction, square the result, and subtract this number from the measured squared error. This procedure is done separately for every lag and segment duration.

We found this procedure substantially reduced the bias when pooling across both random and natural contexts (**Fig S13**, bottom panel), as was done for all of our analyses except those shown in **Figure S4**. When only considering random contexts, we found this procedure somewhat over-corrected the bias (inducing a downward bias for noisy data), perhaps due to the influence of the terms omitted in our approximation (equation 21). However, our results were very similar when using random or natural contexts (**Fig S4**), when using either the uncorrected (**Fig S12**) or bias-corrected error (**Fig 2**), and when using highly denoised data (**Fig S10**). Thus, we conclude that our findings were not substantially influenced by noise and were robust to details of the analysis.

## Acknowledgements

We thank Daniel Maksumov, Nikita Agrawal, Stephanie Montenegro, Leyao Yu, Marcin Leszczynski and Idan Tal for help with data collection. We thank Stephanie Montenegro and Hugh Wang for help in localizing electrodes. And we thank Alex Kell, Stephen David, Josh McDermott, Bevil Conway, Nancy Kanwisher, Nikolaus Kriegeskorte, and Marcin Leszczynski for comments on an earlier draft of this manuscript. This study was supported by the National Institutes of Health (NIDCD-DC014279, S10 OD018211, NINDS-R01-NS084142) and the Howard Hughes Medical Institute (LSRF postdoctoral award to SNH).

---

**Fig S1.**
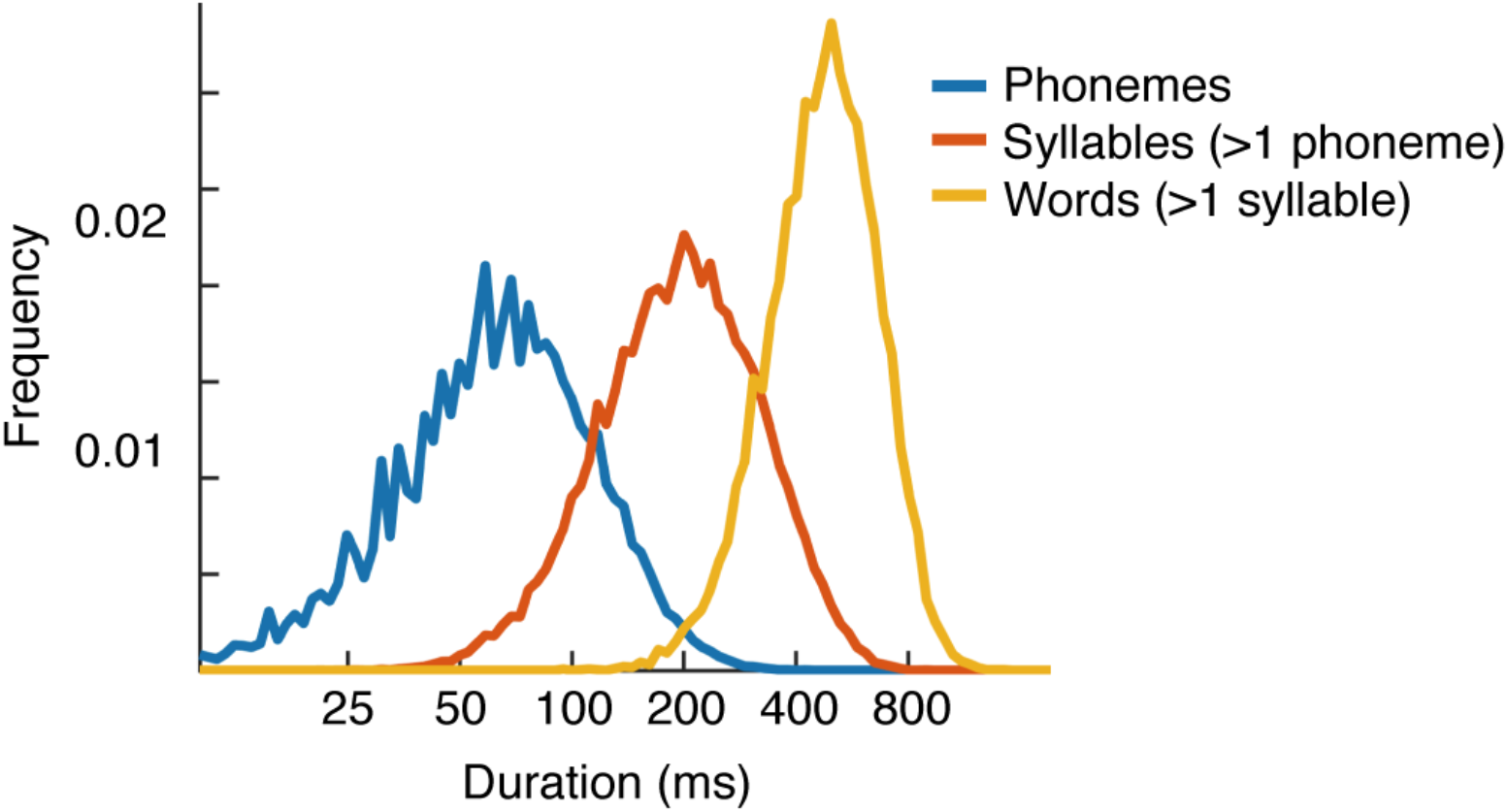
Histogram of phoneme, syllable, and word durations in TIMIT. Durations of phonemes, multi-phoneme syllables, and multi-syllable words in the commonly used TIMIT database. Phonemes and words are labeled in the database. Syllables were computed from the phoneme labels using the software tsylb2^62^. The median duration for each structure is 64, 197, and 479 milliseconds, respectively.

**Fig S2.**
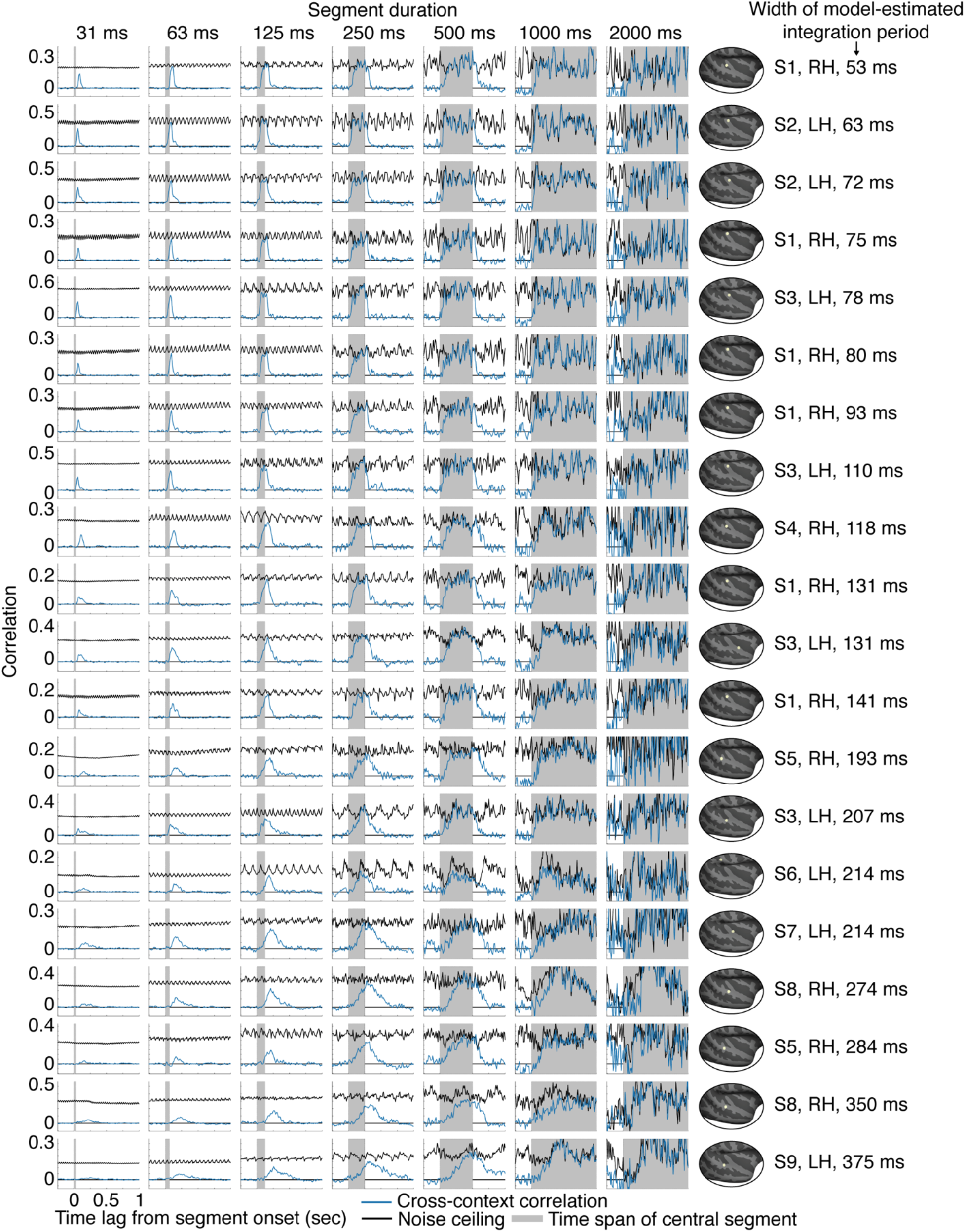
Cross-context correlation for 20 representative electrodes. Electrodes were selected to illustrate the diversity of integration periods. Specifically, we partitioned all sound-responsive electrodes into 5 groups based on the width of their integration period, estimated using a model (**Fig 3** illustrates the model). For each group, we plot the four electrodes with the highest SNR (as measured by the test-retest correlation across the sound set). Electrodes have been sorted by their integration width, which is indicated to the right of each plot, along with the location, hemisphere and subject index for each electrode. Each plot shows the cross-context correlation and noise ceiling for a single electrode and segment duration (indicated above each column). There were more segments for the shorter durations, and as a consequence, the cross-context correlation and noise ceiling were more stable/reliable for shorter segments (the number of segments is inversely proportional to the duration). This property is useful because at the short segment durations, there are a smaller number of relevant time lags, and it is useful if those lags are more reliable. The model used to estimate integration periods pooled across all lags and segment durations, taking into account the reliability of each datapoint (see *Model-estimated integration periods* in the Results and Methods).

**Fig S3.**
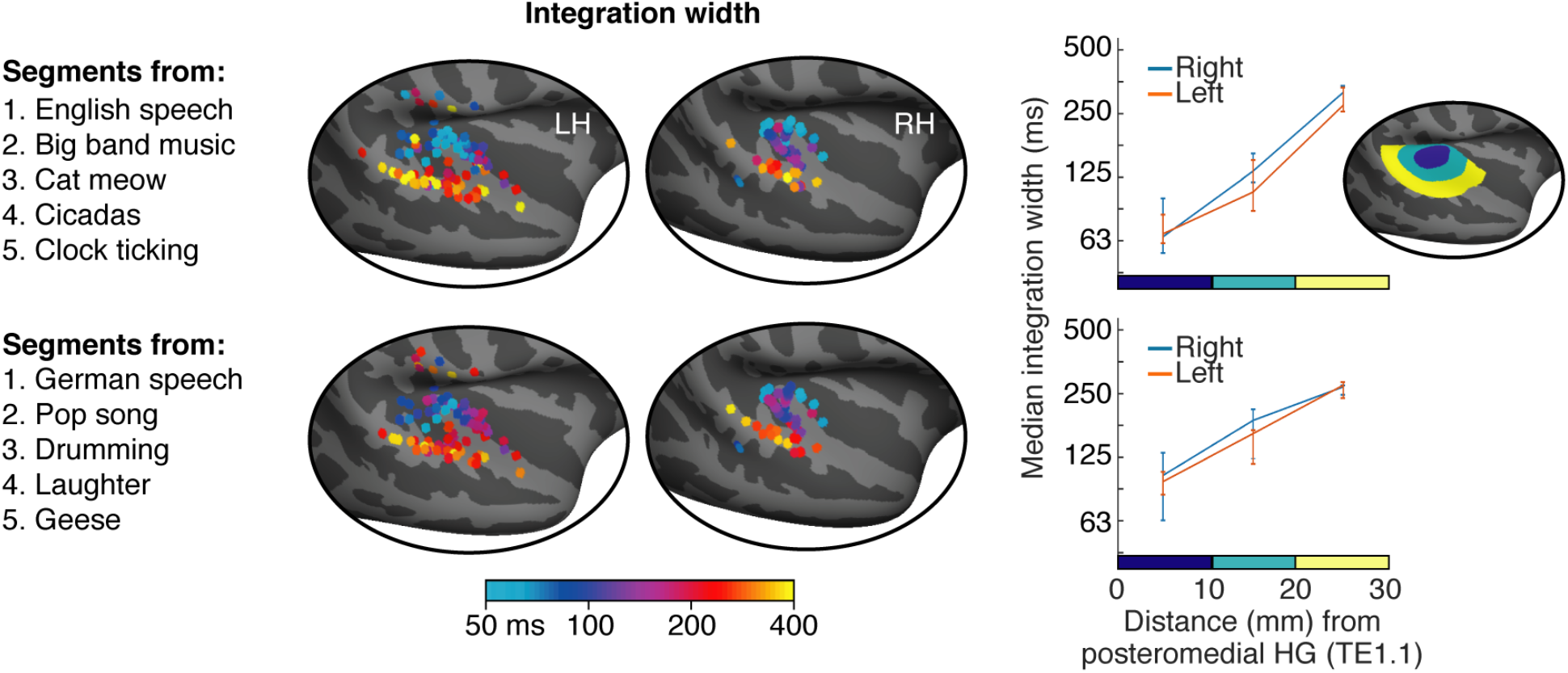
Split-half analysis across the sound set. Sound segments were excerpted from 10 sounds. To assess the robustness of our results to the sounds tested, we estimated integration periods using segments drawn from two non-overlapping splits of 5 sounds each (listed on the left). Since many non-primary regions only respond strongly to speech or music^10–12^, we included speech and music in both splits. Format is analogous to **Figure 4** but only showing integration widths (integration centers were also similar between splits).

**Fig S4.**
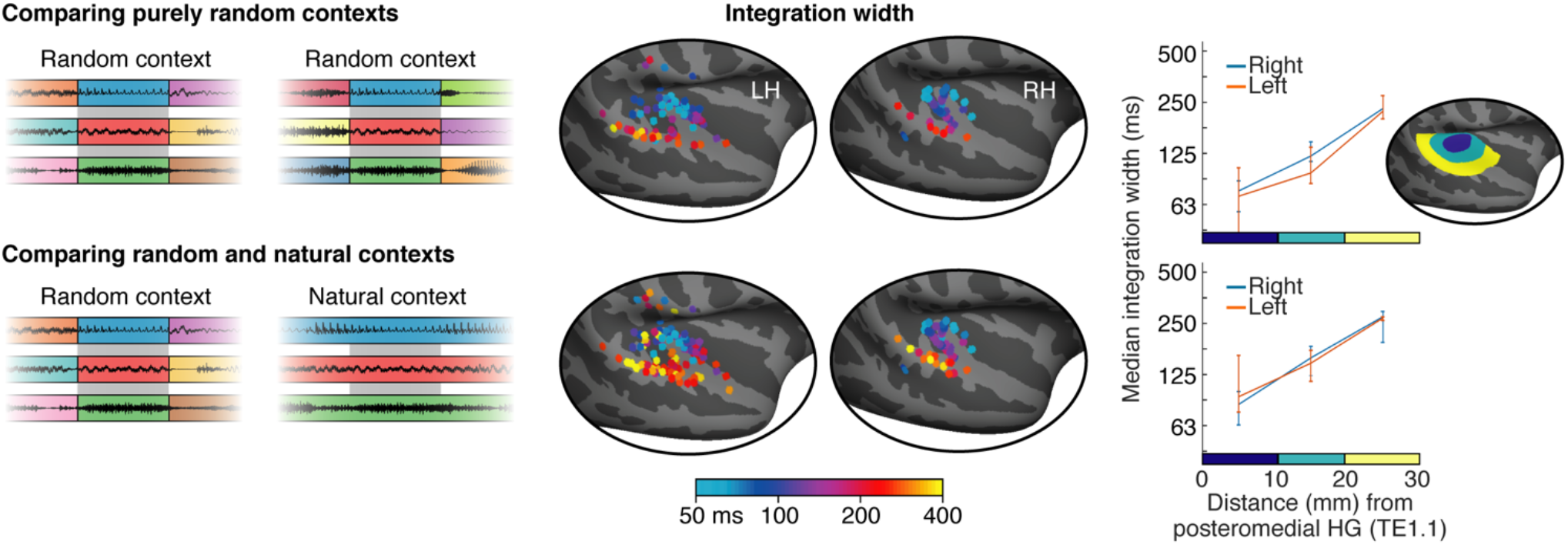
Comparing random and natural contexts. Shorter segments were created by subdividing longer segments, which made it possible to consider two types of context: (1) random context, in which each segment is surrounded by random other segments (2) natural context, where a segment is a subset of a longer segment and thus surrounded by its natural context. When comparing responses across contexts, one of the two contexts must always be random so that the contexts differ. But the other context can be random or natural. Our main analyses pooled across both types of comparison. Here, we show integration widths estimated by comparing either purely random contexts (top panel) or comparing random and natural contexts (bottom panel). Format is analogous to **Figure 4.**

**Fig S5.**
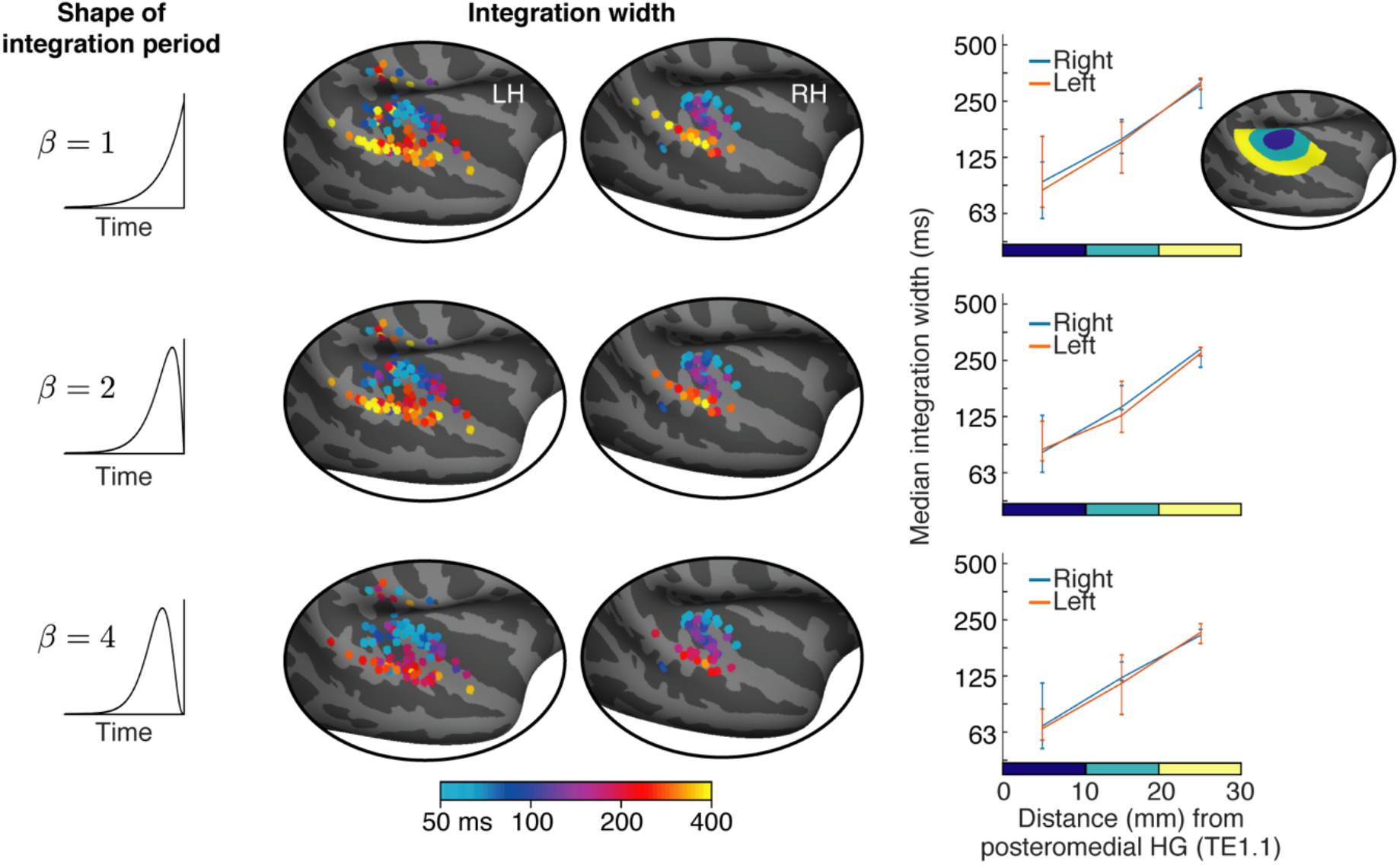
Results for different model window shapes. We modeled integration periods using window shapes that varied from more exponential to more Gaussian (the parameter *β* in equations 1&2 controls the shape of the window, see Methods). For our main analysis, we selected the shape that yielded the best prediction for each electrode. This figure plots integration widths estimated using three different fixed shapes. Format is analogous to **Figure 4.**

**Fig S6.**
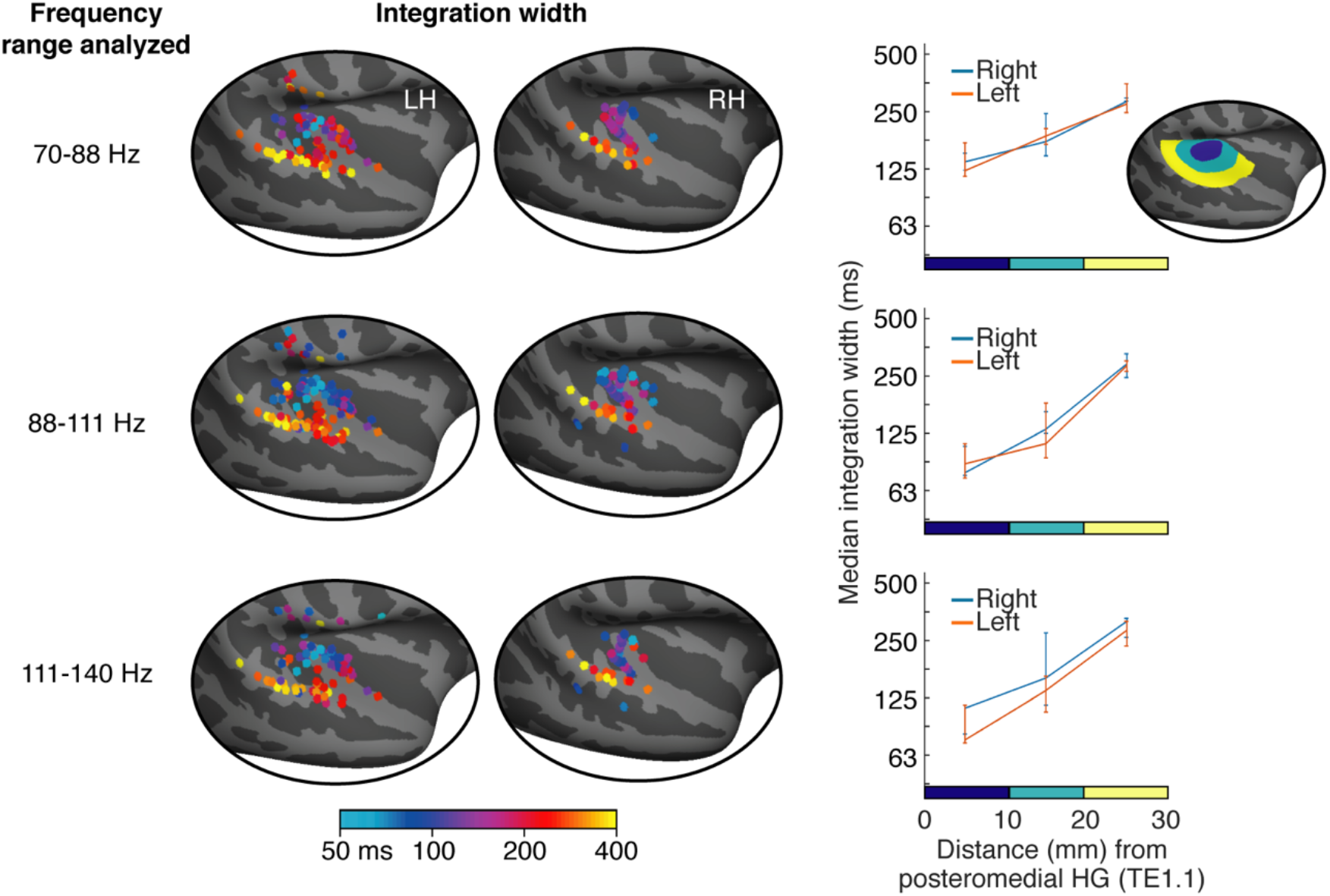
Results for different frequency ranges. For our primary analysis, we measured the broadband power of each electrode between 70 and 140 Hz. This figure shows the results of measuring integration widths from three different subsets of this broader range (equally spaced on a logarithmic scale). Format is analogous to **Figure 4**. Results were similar to those of our main analysis, but using a lower frequency range (70-88 Hz) appeared to limit the shortest integration widths that were detectable by our paradigm, plausibly because faster power fluctuations are better conveyed by a faster carrier.

**Fig S7.**
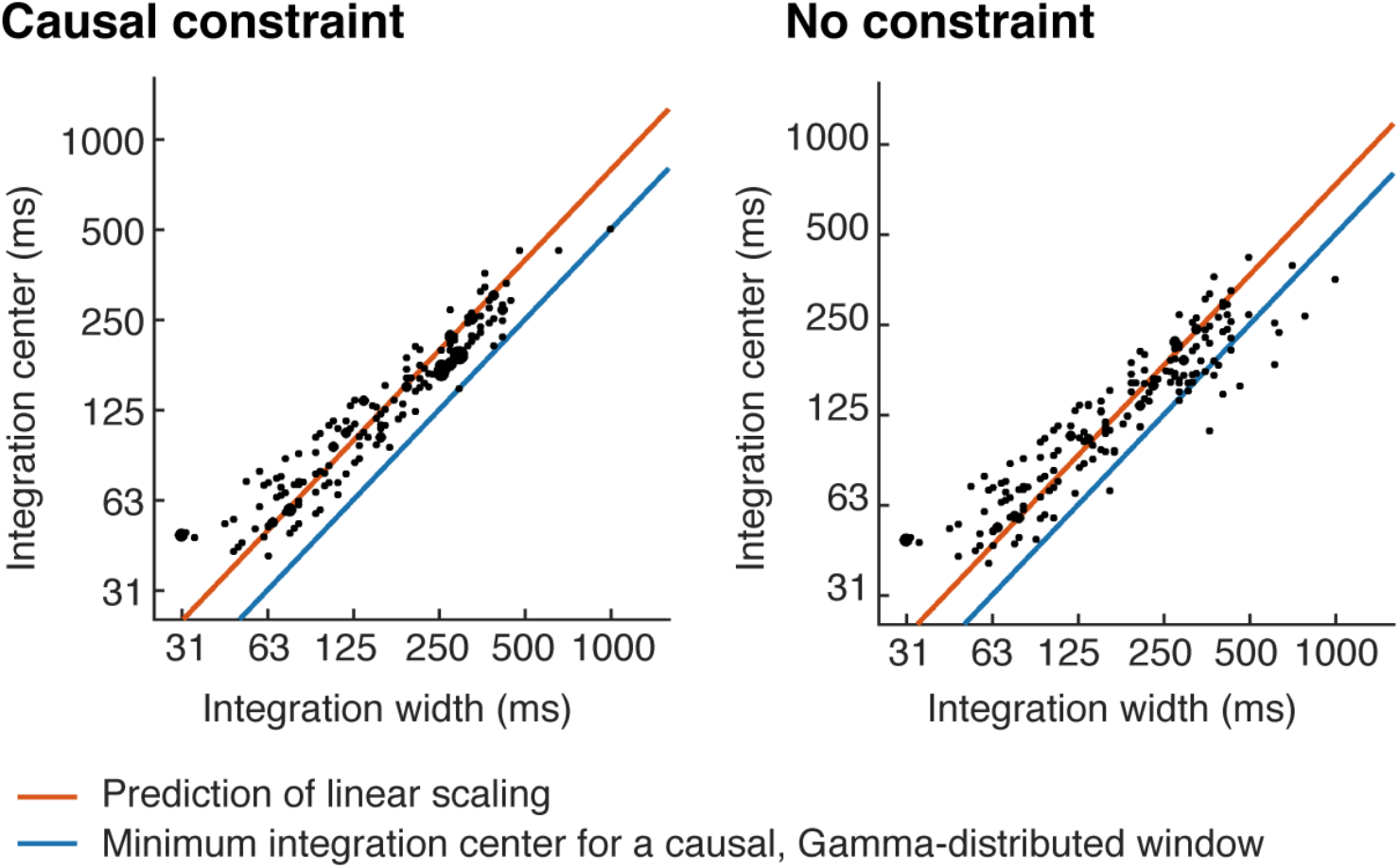
Relationship between integration widths and centers. This figure shows a scatter plot between integration centers and widths. Each dot corresponds to an electrode and larger dots indicate that multiple electrodes were assigned to that pairing of centers/widths. The integration width places a lower bound on the integration center for a causal window (blue line). Integration centers scaled approximately linearly with the integration width (orange line), and remained relatively close to the minimum possible for a casual window. On the left, we show results when integration periods were explicitly constrained to be causal, and on the right, we show results without this constraint. Results were similar in the two cases because the inferred integration periods were close-to-causal without being explicitly constrained to be so.

**Fig S8.**
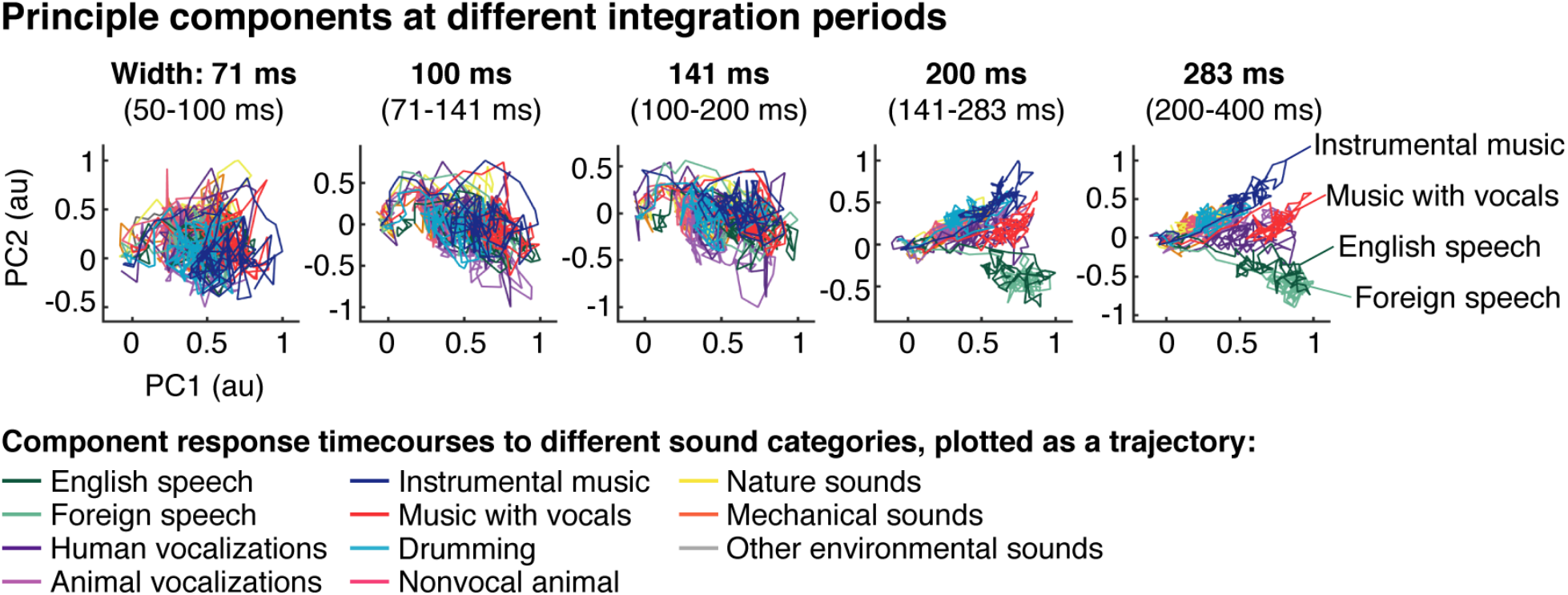
Principal components at different integration periods. Electrodes were grouped based on the width of their integration period in octave intervals (shown above each plot). Responses were then projected onto the top two principal components from each group. This figure shows the average component response timecourse to each category, plotted as a trajectory. Format is the same **Figure 5a**.

**Fig S9.**
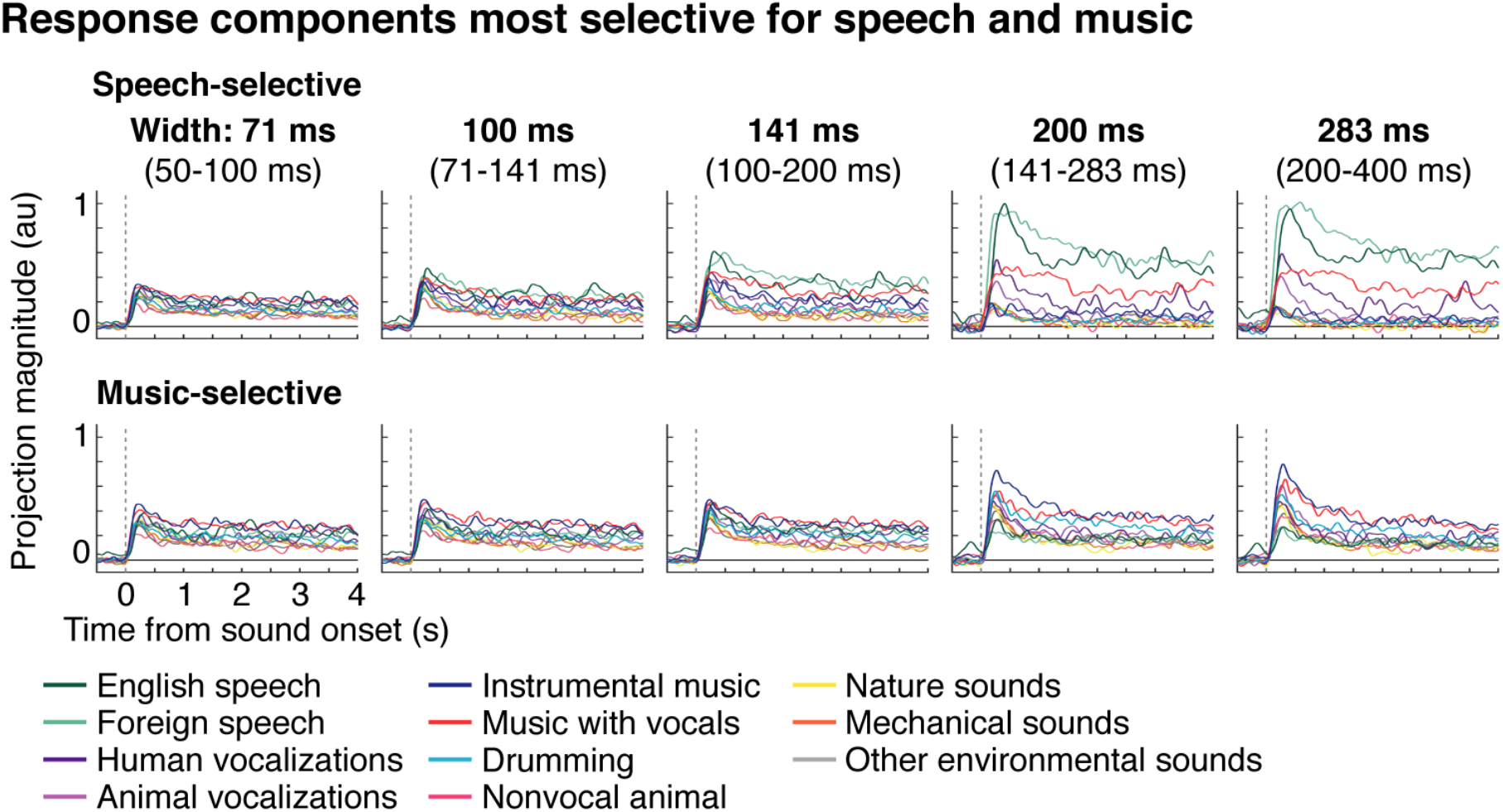
Response components most selective for speech or music. Electrodes were grouped based on the width of their integration period in octave intervals (shown above each plot). The responses from each group were then projected onto the components that showed the greatest speech (top panel) or music selectivity (bottom panel). The speech-selective component was optimized to separate responses to English and foreign speech from all other sounds (excluding vocal music which has speech). The music-selective component was optimized to separate responses to instrumental music, vocal music, and drumming from all other sounds. Each line reflects the average component response timecourse to one sound category. The timecourses have been smoothed so that lines for different categories can be clearly seen (100 millisecond FWHM Gaussian kernel). Independent sounds were used to estimate components and measure their response. **Figure 5c** quantifies how well separated speech and music are along each component. Separation was calculated using the response timecourses to individual sounds without any smoothing (using all timepoints between 250 milliseconds and 4 seconds post-stimulus onset).

**Fig S10.**
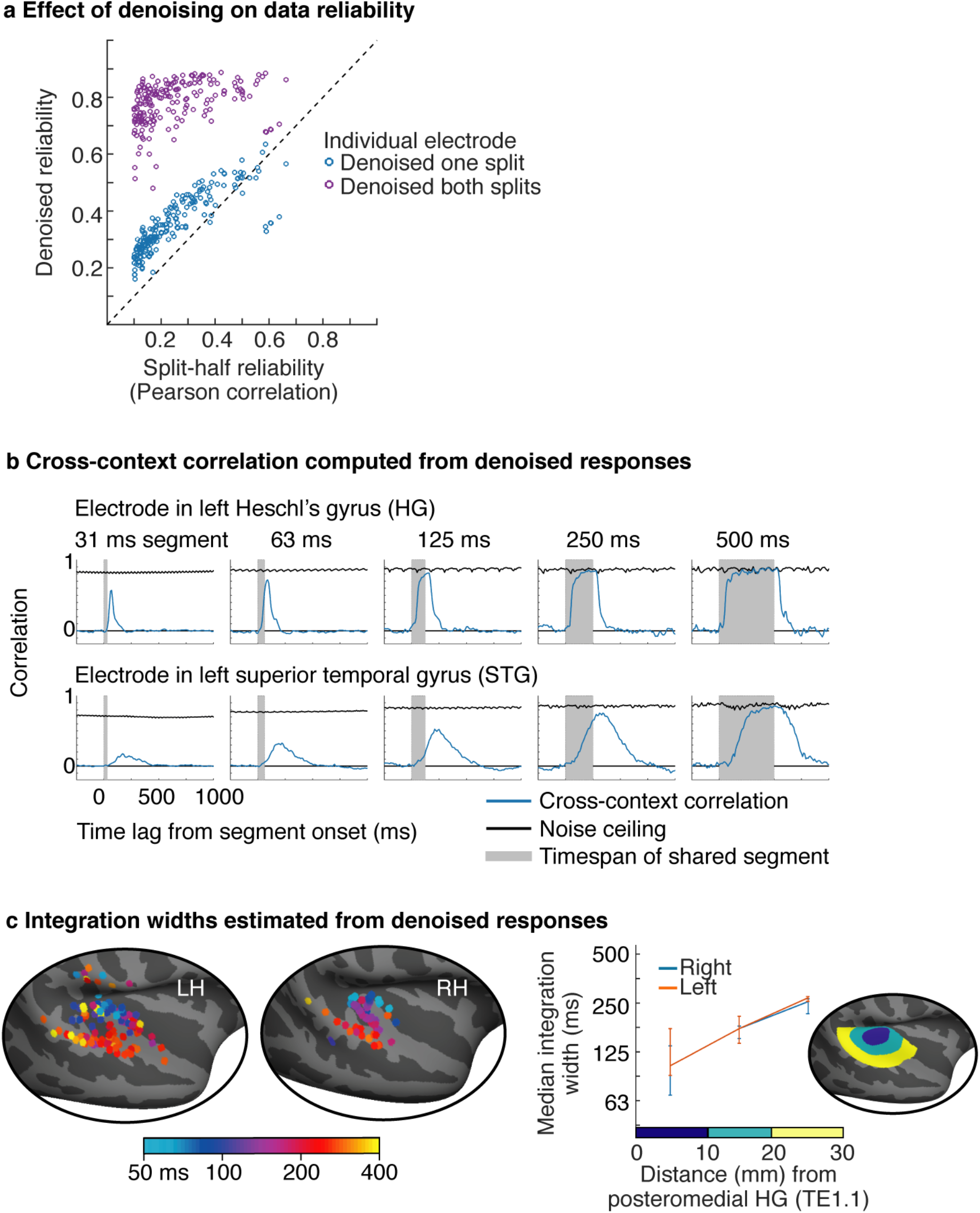
Integration periods estimated from denoised data. **a**, Data were denoised by projecting electrode responses from one subject onto those from all other subjects. This procedure leads to a boost in SNR because the stimulus-driven response is more consistent across subjects than the noise. Each dot shows the split-half reliability of one electrode before (x-axis) or after (y-axis) denoising. The denoising procedure was either applied to both splits of data (purple dots) or to only one split of data (blue dots). Applying the analysis to both splits reveals the overall change in reliability. Applying the analysis to one split provides a fairer test of whether the denoising analysis removes more signal or noise. **b,** The cross-context correlation for the same example electrodes shown in **Fig 2b**, but measured from denoised responses. The trends are similar but the noise ceiling is much higher. **c,** Integration widths estimated from denoised responses. Same format as **Figure 4**.

**Fig S11.**
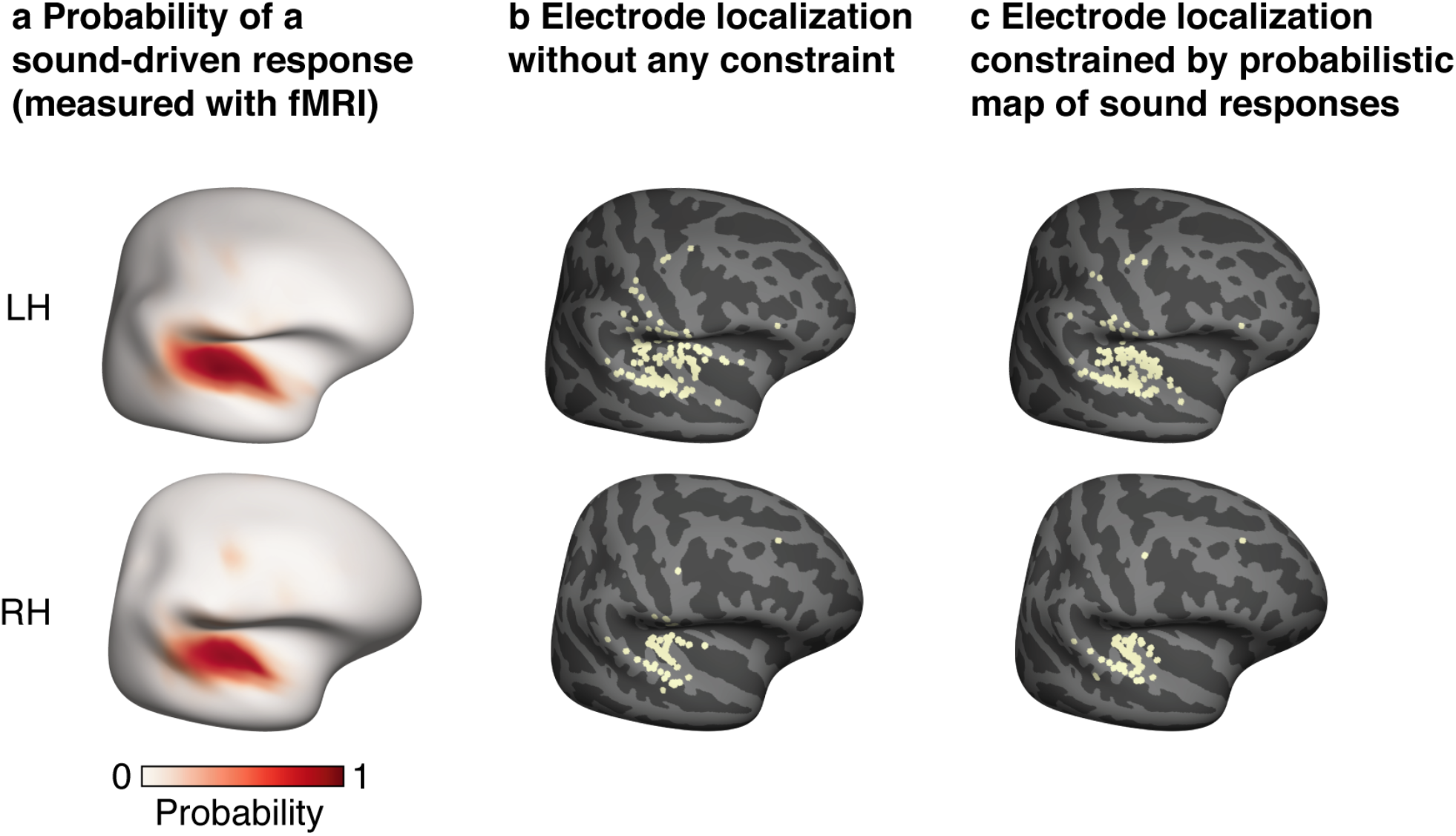
Constraining the anatomical localization of electrodes. **a,** Map showing the probability of observing a significant response to sound at each point in the brain. The map was computed using whole-brain fMRI responses to large collection of 192 natural sounds across a large cohort of 20 subjects^33^. **b,** Electrode localization based on mapping each electrode to the nearest point on the cortical surface. Due to cortical folding, nearby points in space can be faraway on the cortical surface. A s a consequence, small localization errors can cause electrodes to be mapped to the wrong region. Such errors like explain why some electrodes have been localized to the supramarginal gyrus, which abuts the superior temporal gyrus where responses to sound are much more common. **c,** To minimize gross localization errors, we treated the probability map of sound-driven responses shown in panel A as a prior and used to it constrain the localization (see *Electrode localization* in the Methods). Because the prior map is highly smooth this approach did not substantially affect the localization of electrodes at a fine spatial scale.

**Fig S12.**
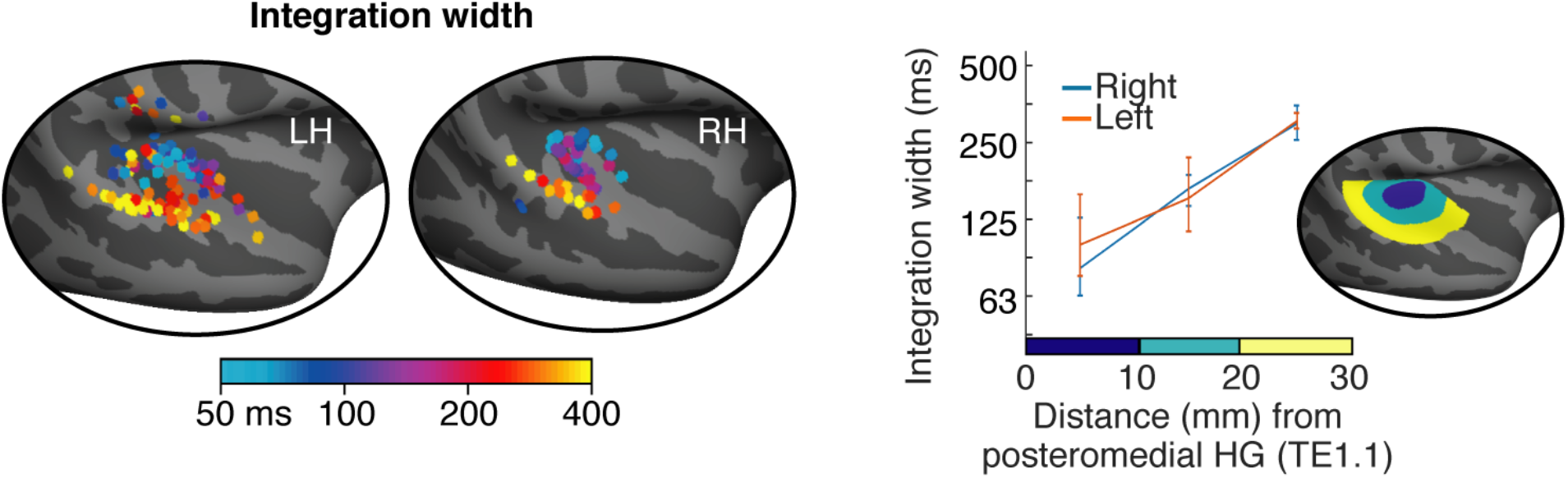
Results using the squared error uncorrected for bias. For our main analysis we quantified prediction accuracy using a bias-corrected variant of the squared error. Here, we plot integration widths estimated using the uncorrected squared error. Results were similar using the uncorrected and bias-corrected error (compare with **Fig 4**), suggesting our data were sufficiently reliable that the bias had little effect.

**Fig S13.**
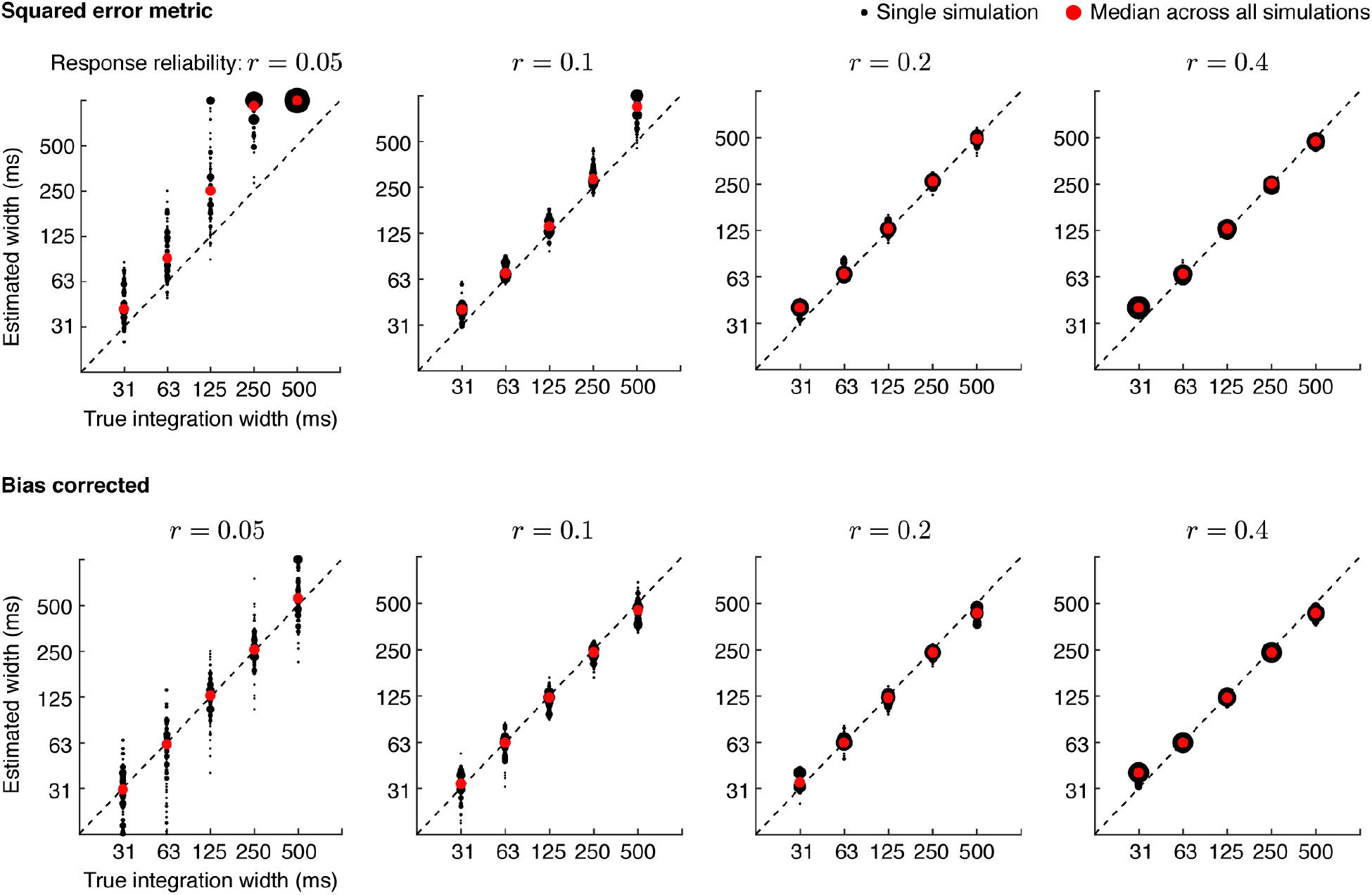
Results of model simulation. We simulated a response that integrated sound amplitude within a Gamma-distributed integration period. The integrated amplitudes were then used to modulate the power of a broadband gamma signal. We tested our ability to infer the correct integration width using our complete analysis pipeline. Black dots show the estimated width for a single simulated response, and red dots show the median width across 100 simulations. Results are plotted for different SNRs, manipulated by adding variable amounts of noise to achieve a desired test-retest reliability (the split-half correlation of the simulated data is shown above each plot). When using the squared error to measure prediction accuracy, we found there was an upward bias for low SNRs (top panel). To address this issue, we derived a variant of the squared error that largely corrected the bias (bottom panel) (see *Bias correction* in Methods).

**Table S1.**
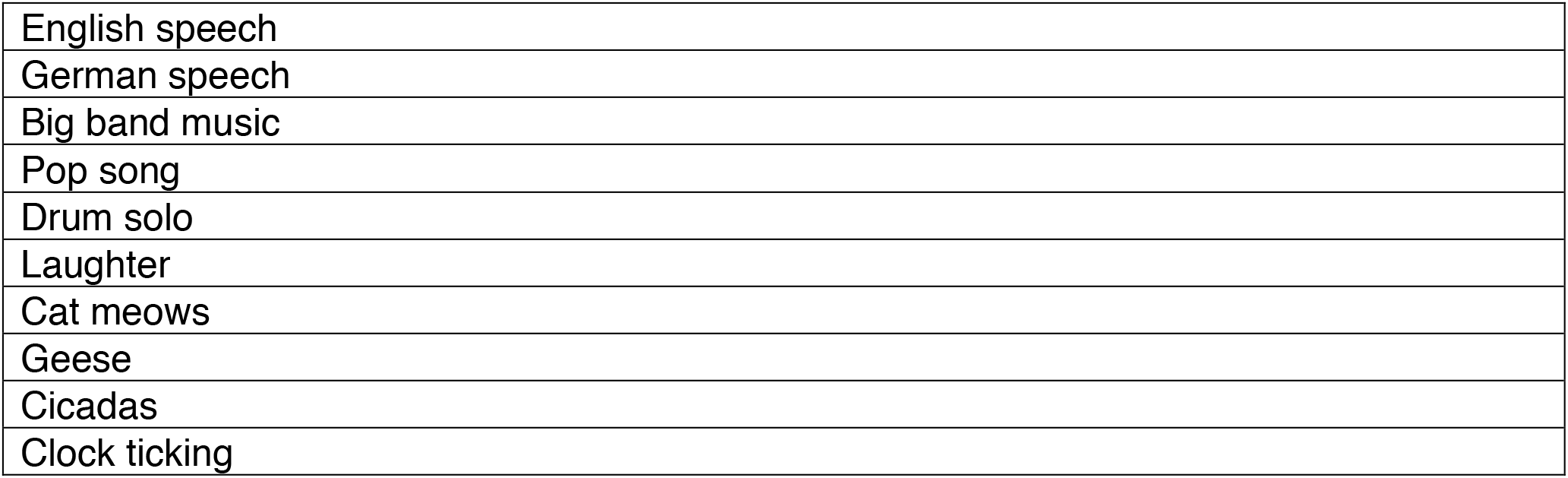
Sound sources used for TCI experiment.

